# Contributions of adaptation and purifying selection to SARS-CoV-2 evolution

**DOI:** 10.1101/2022.08.22.504731

**Authors:** Richard A. Neher

**Affiliations:** Biozentrum, University of Basel, Basel, Switzerland; Swiss Institute of Bioinformatics, Switzerland

## Abstract

Continued evolution and adaptation of SARS-CoV-2 has lead to more transmissible and immune-evasive variants with profound impact on the course of the pandemic. Here I analyze the evolution of the virus over 2.5 years since its emergence and estimate rates of evolution for synonymous and non-synonymous changes separately for evolution within clades – well defined mono-phyletic groups with gradual evolution – and for the pandemic overall. The rate of synonymous mutations is found to be around 6 changes per year. Synonymous rates within variants vary little from variant to variant and are compatible with the overall rate of 7 changes per year (or 7.5 × 10^−4^ per year and codon). In contrast, the rate at which variants accumulate amino acid changes (non-synonymous mutation) was initially around 12-16 changes per year, but in 2021 and 2022 dropped to 6-9 changes per year. The overall rate of non-synonymous evolution, that is across variants, is estimated to be about 26 amino acid changes per year (or 2.7 × 10^−3^ per year and codon). This strong acceleration of the overall rate compared to within clade evolution indicates that the evolutionary process that gave rise to the different variants is qualitatively different from that in typical transmission chains and likely dominated by adaptive evolution. I further quantify the spectrum of mutations and purifying selection in different SARS-CoV-2 proteins and show that the massive global sampling of SARS-CoV-2 is sufficient to estimate site specific fitness costs across the entire genome. Many accessory proteins evolve under limited evolutionary constraint with little short term purifying selection. About half of the mutations in other proteins are strongly deleterious.

---

Since its emergence in late 2019 (Zhu *et al*., 2020), SARS-CoV-2 has displayed a discontinuous pattern of evolution with large jumps in sequence space giving rise to phylogenetically distinct variants (Faria *et al*., 2021; Hodcroft *et al*., 2021; Naveca *et al*., 2021; Tegally *et al*., 2021; Viana *et al*., 2022; Volz *et al*., 2021). Many of these variants spread considerably faster and quickly displaced the resident variants at the time either because of intrinsically increased transmissibility, evasion of prior immunity in the population, or a combination of both. Specific Variants of Concern (VoC) or Interest (VoI) were designated by the WHO and labeled by Greek letters (Konings *et al*., 2021). The branches leading to these variants are characterized by many amino acid-changing mutations that often cluster in the S1 domain of the spike protein (Kistler *et al*., 2022).

This pattern of rapid non-synonymous evolution in viral surface proteins that interact with the host cells is common among many RNA viruses and for example well studied in influenza A virus evolution (Bhatt *et al*., 2011; Strelkowa and Lässig, 2012). But adaptive evolution of influenza viruses tends to be gradual without large jumps in sequence space, while new variants of SARS-CoV-2 with tens of novel mutations emerged suddenly without intermediate genomes being observed, the most dramatic being the emergence of Omicron in late 2021 (Viana *et al*., 2022). The pre-dominant hypothesis for this cryptic emergence of highly mutated, transmissible and immune evasive variants are chronic infections that are common in patients with impaired immune systems, either through HIV-1 infection (Cele *et al*., 2022) or medical intervention (Choi *et al*., 2020; Kemp *et al*., 2021). During such chronic infections, extensive intra-host diversity can develop through accelerated evolution (**?**). Onward transmission from such chronic infections has also been documented (Gonzalez-Reiche *et al*., 2022). However, to date there is no direct evidence for the mode of emergence of any variant. The case for chronic infection being an important contributor is strongest for the variants Alpha and Omicron (Hill *et al*., 2022).

The dichotomous pattern of SARS-CoV-2 evolution with step-wise evolution within variants and atypical bursts of evolution leading to new variants has been investigated by Tay *et al*. (2022) and Hill *et al*. (2022), who showed that the rate of evolution along branches giving rise to new variants is up to four-fold higher than the background rate. Here, I build on these results and investigate the patterns of SARS-CoV-2 diversification within variants and compare these to the global dynamics of evolution and adaptation. This comparison reveals a consistent dichotomy between slow within-variant evolution and rapid adaptive evolution giving rise to new variants. This difference in evolutionary rate is only seen for non-synonymous changes – the rate of synonymous evolution within variants is compatible with that seen between variants. Furthermore, early variants display more rapid non-synonymous evolution than later variants suggesting more ubiquitous adaptive evolution early on. I further quantify the level of functional constraint of different open reading frames and infer a map of mutational tolerance across the genome from patterns of rare diversity.

## Results

Evolutionary rates and divergence times are typically estimated using phylogenetic approaches (Drummond *et al*., 2006). These methods, however, cannot handle the volume of SARS-CoV-2 data available and data have to be dramatically down-sampled. Furthermore, phylogenetic methods impose an hierarchical structure on the data and are thus very sensitive to problematic sequences or metadata: Any mis-dated, recombinant, contaminated, or otherwise chimeric sequence can fatally distort the analysis. Problematic sequences can be particularly common when a new variant takes over since sequencing protocols need adjusting (De Maio *et al*., 2020).

To circumvent many of the above-mentioned problems and still use the majority of the available data, I use a combination of automated filtering and simple robust approaches to analyze the evolutionary patterns (see Materials and Methods). I first use Nextclade (Aksamentov *et al*., 2021) to assign 12 million sequences available in GISAID (2022-07-25) (Shu and McCauley, 2017) to one of the Nextstrain clades (Hadfield *et al*., 2018; **?**), which are denoted by a Year-Letter combination (e.g. 19A, 20A, 20B, …), see Fig. 1. Sequences belonging to recombinant Pango lineages are excluded (Rambaut *et al*., 2020). These clades represent well-defined groups of sequences with little evidence of recombination within them and are analyzed independently. In this analysis, I only consider clades that have had significant circulation for at least 6 months.

**FIG. 1.**
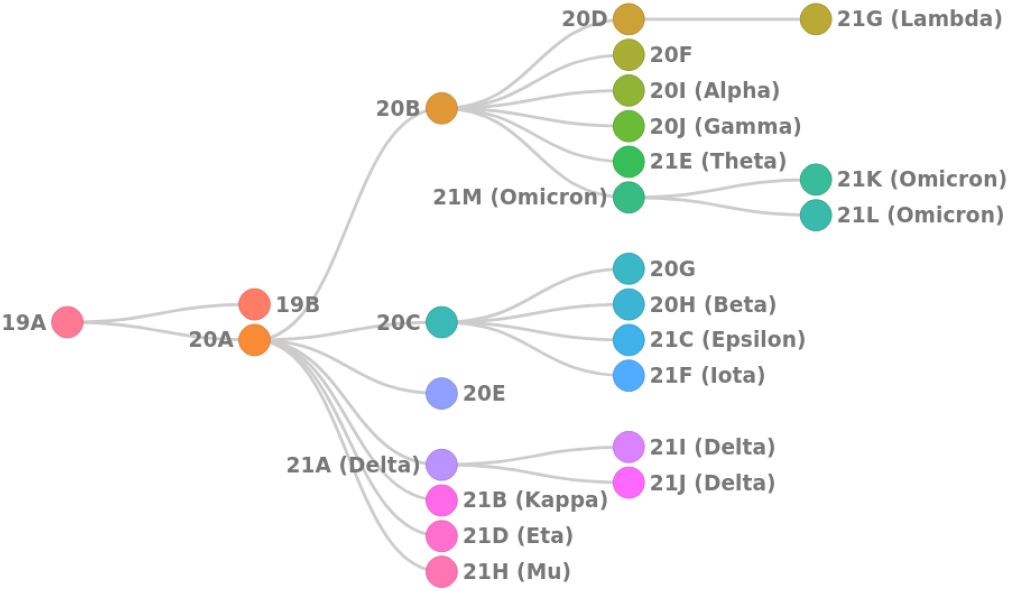
Schematic summary of Nextstrain clades.

For each clade, I define a “founder” genotype and exclude any sequence that does not have the full set of clade defining mutations relative to the reference sequence Wuhan/Hu-1. This founder sequence is manually curated for each clade considered. This filtering removes most incomplete sequences as well as sequences where amplicon drop-out are back-filled with the reference sequence, but will ignore a few sequences with true reversion mutations. In addition, sequences with a Nextclade QC score above 30 (80 for 21H because of an unaccounted frameshift) are removed. Results are insensitive to the stringency of this filtering. For this reduced set of sequences, I determine the mutations they carry on top of the founder genotype of the clade to analyze diversification and divergence within the clade. The latter step is done for nucleotide changes as well as for amino acid changes.

Within each clade, the number of mutations is expected to increase linearly in time and the variation around this mean would, in an ideal case, obey Poisson statistics. For the majority of sequences, this is approximately true, but some problematic sequences have more mutations than expected. To exclude these outliers, I perform a simple linear regression of the number of “intra-clade” mutations against time and remove sequences whose deviation from the linear fit exceeds twice the expected standard deviation by 3 mutations (see Fig. 2). These outliers are never more than 1% of all sequences that pass the first round of filtering and the results are insensitive to the filtering criteria.

**FIG. 2.**
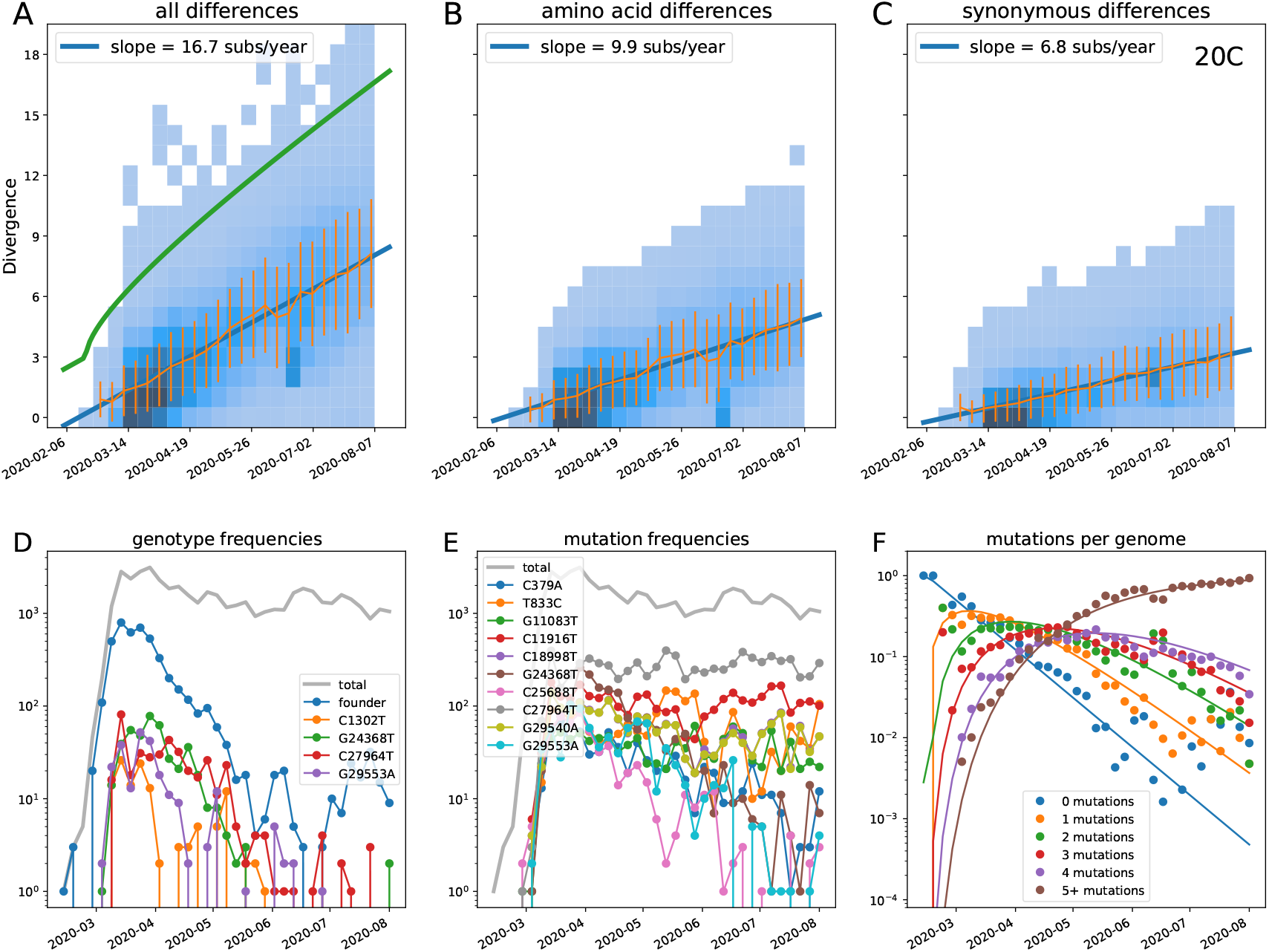
Within-clade divergence increases linearly with time (example clade 20C). Top: Each panel shows the number of within-clade mutations (total (A), amino acid changing (B), synonymous (C)) as a function of time. The green line in panel A indicates the divergence cut-off, panels B&C only show sequences that pass the divergence filter. Each panel also shows mean *±* standard deviation and a weighted linear fit. Bottom: Panels D and E show the prevalence of the most common genotypes (D) and mutations (E) during the first 3 months. Genotypes in (D) refer to sequences with **all and only** the mutations indicated, while mutations in (E) count all sequences with a specific mutation, regardless of mutations elsewhere. In the case shown (20C), the founder genotype initially dominates and no specific mutation ever reaches high frequency. Note that specific genotypes invariably decrease in frequency due to subsequent mutations, while mutations irrespective of their genetic background persist. Panel F shows a Poisson model fit to the breakdown of the population into genotypes with different number of mutations over time. Analogous figures for other clades are included in the appendix.

After removing these outliers, the data are binned by calendar week. Evolutionary rate and putative emergence date of the variant are then estimated by weighted linear regression where each time bin is weighted with the fourth root of the number of sequences in the bins. The exact functional form of this weighting does not have a big influence on the results, but a sublinear weighting helps to counter the large variation in sequencing effort across countries and the unavoidable imbalance due to the fact that few sequences are available early on when an emerging variant is still rare. Fig. 2 shows the increasing intra-clade divergence for clade 20C, a large clade that emerged in early 2020 that was common in North America and Scandinavia. Both synonymous and non-synonymous within-clade average divergence increase linearly over time allowing for a robust estimate of the rate.

Due to shared ancestry, divergences of sequences are not independent data points and a regression against time is general not a suitable method to estimate evolutionary rates. In particular, confidence intervals are difficult to obtain. However, in the case of rapidly expanding variants we typically observe a large number of independent lineages emanating from one or several basal polytomies. Along each of these lineages, mutation accumulation is independent. While not every sequence is an independent sample, the effective number of independent samples is large and the steadily increasing average divergence allows to estimate the rate robustly.

A simple model for diversity within a growing variant is a super-critical branching process with growth rate *α* and an embedded mutation process. Offspring of genomes with *i* mutations will carry *i* + *j* mutations, where *j* is a Poisson distributed number with mean *μt* (mutation rate *μ* and generation time *t*). The probability that offspring genomes are different from their parents is *u* = 1 − *e*^−*μt*^, which for a generation time of *t* = 5 days and a rate of *μ* = 15*/year* evaluates to *u* ≈ 0.2 (note that this rate excludes strongly deleterious mutations, see Discussion).

When considering a rapidly growing well sampled outbreak, typically a single founder genotype will give rise to a large number of daughter lineages that evolve independently. In this case, the diversification processes is robustly described by its mean. Since the above branching process is linear, the mean number of cases *n* will increase exponentially with rate *α*, while the number of genomes with *i* mutations relative to the founder *m*_*i*_ grows with rate *α* − *u* per generation:

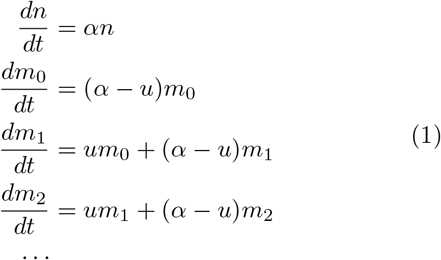

with solution 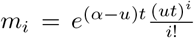. Note that this model assumes continuous time and only allows increments by one mutation at a time. Multiple mutations in one serial interval can still happen through successive mutations within one host. At time *t* after the emergence of the variant, the number of mutations in the population is expected to be Poisson distributed with mean *ut*. The overall number of cases is *e*^*αt*^ or more generally _*e*_ *∫*^*α*(*t*)*dt*^ if growth rate or ascertainment varies over time.

For some clades, especially those that are well sampled soon after their emergence, this Poisson model is a good fit to diversity accumulation and yields estimates of rates and time of origin that are compatible with the divergence regression, see Fig. 2F for clade 20C. In this case, the founder genotype initially dominates, but is gradually replaced – first by single mutant genotypes, then double mutants, and so forth (comp. Fig. 2). Analogous graphs for all other clades considered are included in the appendix.

In other cases, notable subclades did become dominant during the early stochastic dynamic. In this case, extrapolation of the linear fit to zero divergence is not necessarily a good estimator of the emergence time of a variant: if branches leading to big subclades carry anomalously many or few mutations, the divergence time will be over- or underestimated. For several clades 19B, 20H (Beta), 21D (Eta), 21G (Lambda), 21H (Mu), 21I (Delta), 21J (Delta), 21L (Omicron, BA.2), and 22B (Omicron, BA.5), founder-like variants are a minority even in early data.

These Poisson weights of mutation numbers are again only valid if mutations accumulate along many independent lineages. In particular, this assumption is violated if some lineages spread systematically faster than others either because of epidemiological factors or because they carry adaptive mutations. In variant 21K (Omicron, BA.1), a sublineage with mutation S:R346K might have enjoyed a transmission advantage. A possible consequence of this advantage is visible in Supp. Fig. 21 where genotypes with 0, 1, or 2 mutations are decaying more rapidly than those with more mutations.

Despite these caveats, for almost all Nextstrain clades, the slope at which diversity increases is robust (similar patterns as Fig. 2A-C for clade 20C), allowing us to estimate clade-specific evolutionary rates for amino acid and synonymous changes. These rates are summarized in Fig. 3 and Tab. I. In addition to these within-clade rates, I estimated the rate at which the clades themselves accumulated amino acid and synonymous changes by regressing the number of differences of the clade’s founder sequence (relative to the putative root in clade 19B (Caraballo-Ortiz *et al*., 2022)) against the estimated time of origin of the clade. These regressions are shown in thick gray lines in Fig. 3.

**TABLE I.** Evolutionary rates estimates from root-to-tip regressions for overall nucleotide changes, amino acid changes, and synonymous changes. The column ‘# of seq.’ shows the number of sequences that entered the analysis after filtering. The last three columns give the distances of the clade founder sequence from putative MRCA of SARS-CoV-2 (19B).

| clade | # of seq. | overall rate [$y^{-1}$] | aa rate [$y^{-1}$] | syn rate [$y^{-1}$] | overall div. | aa div. | syn div. |
| --- | --- | --- | --- | --- | --- | --- | --- |
| 19B | 8187 | 19.46 | 12.37 | 7.09 | 0 | 0 | 0 |
| 19B+ | 22745 | 18.56 | 11.92 | 6.64 | 0 | 0 | 0 |
| 19B++ | 126823 | 27.91 | 16.41 | 11.50 | 0 | 0 | 0 |
| 19A | 14382 | 19.73 | 12.79 | 6.95 | 2 | 1 | 1 |
| 20A | 44629 | 15.08 | 7.94 | 7.15 | 6 | 3 | 3 |
| 20B | 42077 | 14.29 | 7.69 | 6.59 | 9 | 5 | 4 |
| 20C | 47865 | 16.73 | 9.92 | 6.81 | 8 | 5 | 3 |
| 20A+ | 136863 | 16.30 | 9.40 | 6.90 | 6 | 3 | 3 |
| 20E | 82240 | 14.04 | 5.65 | 8.39 | 13 | 5 | 8 |
| 20H (Beta) | 5077 | 15.48 | 9.51 | 5.96 | 21 | 16 | 5 |
| 20I (Alpha) | 277226 | 11.56 | 5.49 | 6.07 | 30 | 19 | 11 |
| 20J (Gamma) | 24970 | 12.97 | 6.29 | 6.68 | 35 | 22 | 13 |
| 21D (Eta) | 3653 | 11.36 | 6.29 | 5.07 | 27 | 13 | 14 |
| 21G (Lambda) | 4567 | 12.04 | 6.51 | 5.53 | 28 | 19 | 9 |
| 21H (Mu) | 2897 | 18.25 | 10.05 | 8.20 | 31 | 21 | 10 |
| 21I (Delta) | 86585 | 16.55 | 8.93 | 7.62 | 30 | 24 | 6 |
| 21J (Delta) | 733825 | 14.30 | 7.17 | 7.14 | 34 | 28 | 6 |
| 21K (Omicron) | 956696 | 13.48 | 6.45 | 7.03 | 56 | 46 | 10 |
| 21L (Omicron) | 690177 | 9.24 | 4.39 | 4.85 | 68 | 50 | 18 |
| 22A (Omicron) | 33451 | 7.71 | 2.81 | 4.90 | 71 | 52 | 19 |
| 22B (Omicron) | 124239 | 11.90 | 5.03 | 6.87 | 66 | 50 | 16 |

**FIG. 3.**
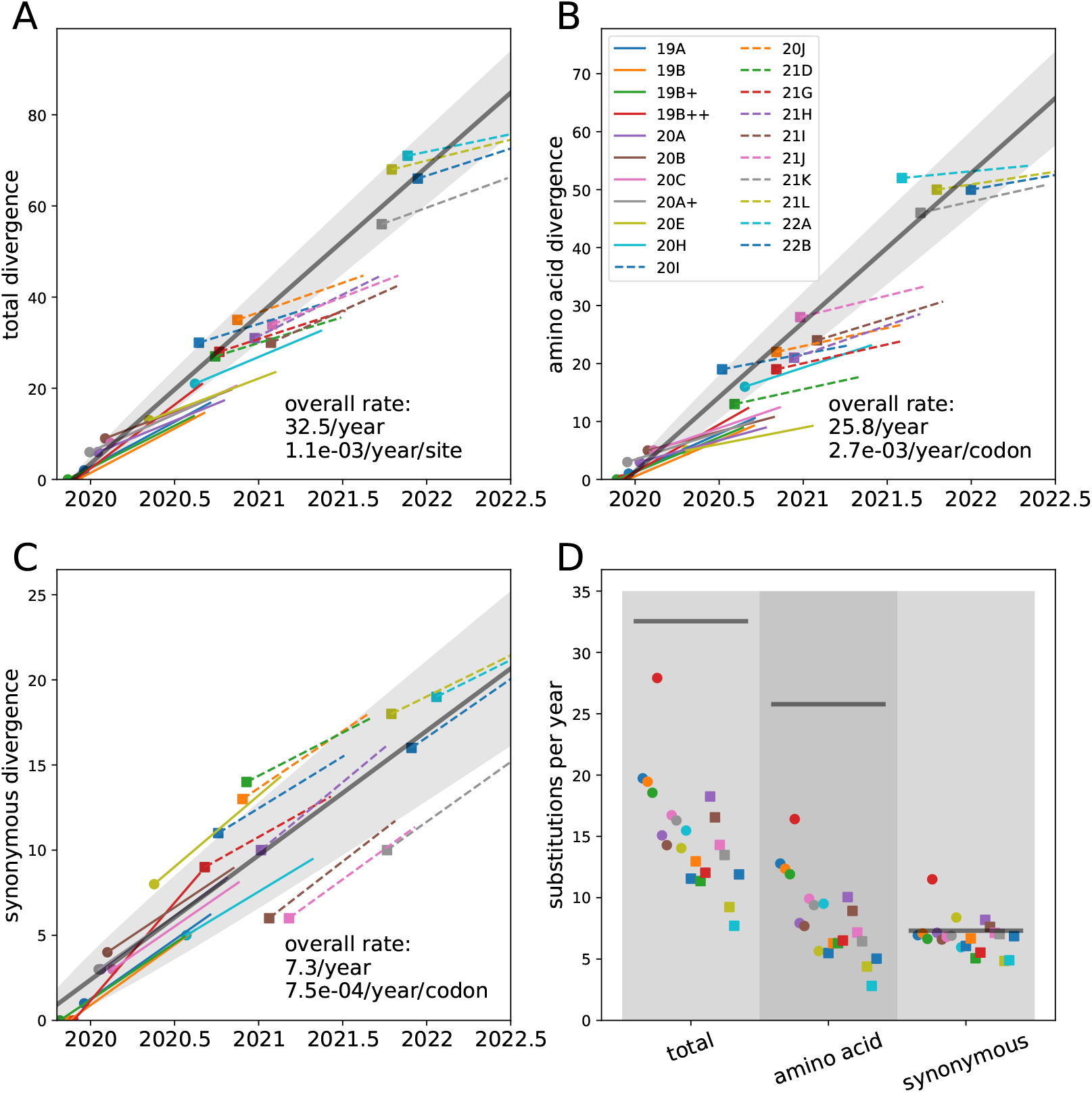
Divergence and evolutionary rates of different Nextstrain clades. Panels A,B&C show the estimated divergence of the founder genotype of each clade (big dot or square) and the subsequent divergence trend for all nucleotide changes, amino acid changes, and synonymous changes, respectively. In addition, each panel contains a regression of the divergence of clade founders vs time (gray line). The standard deviation expected based on Poisson statistics is indicated as shaded area. Panel D summarizes the individual rate estimates (dots and squares) and compares them to the estimate of inter-clade rates (gray lines). In panel D, clades are in alphabetical order, which is similar to their (uncertain) order of emergence. The red outlier is ‘composite’ clade 19B++ (containing 19B, 19A, 20A, 20B, 20C) with inflated rates due to adaptive mutations on the branch leading to clade 20A.

Rates of synonymous change are very consistent across clades (about 5-8 changes per genome per year) and also agree with the overall rate of synonymous changes of 7.3 changes per genome per year. The rates of non-synonymous changes are much more variable (Fig. 3D). Within clades, the rate of non-synonymous changes varies between 5 and 16 changes per year. Earlier clades are estimated to have larger rates around 10-15 changes per year, while rate estimates for later clade fall between 3 to 9 changes per year (see Fig. 4). In contrast, the interclade non-synonymous rate exceeds 25 changes per year. The spike protein does not contribute to this decreasing trend in the rate of within-clade non-synonymous evolution, which is most evident in *ORF1ab* and to a lesser degree in accessory proteins and *N* (see Supp.Fig. 1).

**FIG. 4.**
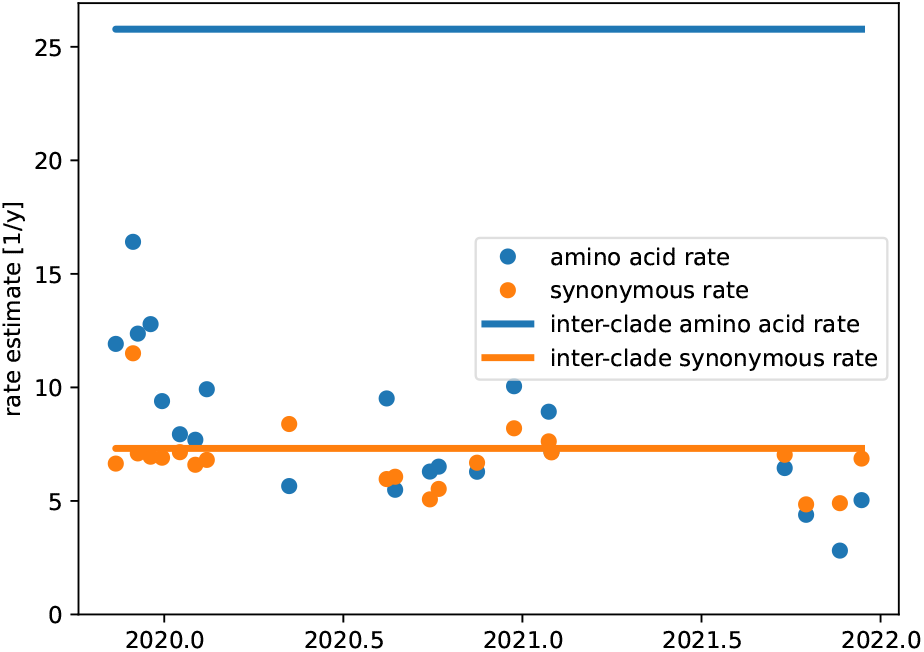
Divergence and evolutionary rates of different Nextstrain clades over time. Synonymous rates estimates are stable in time and fluctuate around the rate estimates for between clades. Non-synonymous rate estimates are highest for clades 19A to 20C that emerged early in the pandemic.

Nextstrain clades tend to be defined by long branches leading to a large polytomy. It could thus be that the estimated inter-clade rate exceeds the intra-clade rate purely because of this conditioning. This effect might be particularly important early on in the pandemic when diversity was low and when branches with as few as two mutations were used to define new clades. I therefore also included composite clades 19B+, 19B++ and 20A+ containing sequences from 19A and 19B (rooted on 19B), 19A, 19B, 20A, 20B, 20C, 20D (rooted on 19B), and 20A, 20B, 20C, 20D (rooted on 20A). The estimates for composite clades 19B+ and 20A+ are consistent with the estimates of the individual clades, while apparent rates of 19B++ are considerably higher. The latter is due to the rapid expansion and subsequent dominance of clade 20A, the first clade with spike mutation *D*614*G*, and its descendants which rapidly fixed four additional mutations (Korber *et al*., 2020). This is an early example of an accelerated global rate of evolution due to adaptive evolution, see Discussion.

At the other extreme, some Omicron clades have very low rate estimates which should be interpreted carefully. Founder genotypes of Omicron clades 21L (BA.2), 22A (BA.4) and 22B (BA.5) are rarely sampled. These clades showed considerable diversity, including reversions to the reference, soon after their initial discovery (Tegally *et al*., 2022). This diversity and the complex, possibly recombinant, origin of the Omicron clades makes estimating their rates of evolution challenging.

### Purifying selection and mutation tolerance

As described above, the rate of synonymous mutations is comparable within and between clades without a strong indication that this rate might have changed over time. This is expected, as synonymous positions are rarely a locus of adaptation and tend to have a small effects on fitness in large parts of the genomes of RNA viruses (Zanini *et al*., 2017) (outside of specific regions with important RNA elements or splice sites). To quantify how much of the SARS-CoV-2 genome is constrained and how strongly purifying selection operates on different genomic regions, I made use of the “rare mutations” annotation provided by Nextclade. Nextclade attaches each sequence to a reference tree and determines by which mutations it differs from the attachment point. For each Pango lineage (as determined by Nextclade) (Aksamentov *et al*., 2021; Rambaut *et al*., 2020), I count how often these “rare mutations” (including reversions to the reference) are observed. This way, for each position in the genome, one obtains the fraction of lineages with minor variation (excluding singletons). I normalize this fraction against the relative rate of mutation away from the ancestral nucleotide (see Supp. Fig. 2), and use this as a semi-quantitative proxy of mutational tolerance.

Simply splitting the genome into at 1st, 2nd, and 3rd positions of codons already reveals strong signatures of purifying selection, see Fig. 5. Between 15 and 20% of first and second positions in codons show almost no variation, while half of these sites are less variable than the most constrained 10% of third positions. The median of variation at 3rd positions is more than double that at 1st and 2nd positions.

**FIG. 5.**
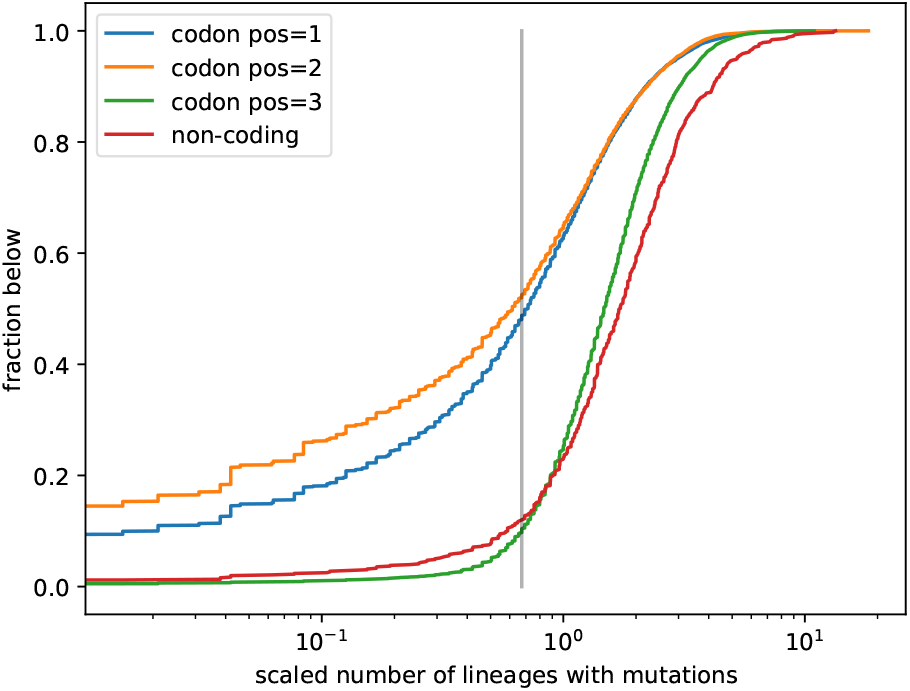
Constraints on SARS-CoV-2 mutations. Almost all 3rd codon positions tolerate mutations, while 1st and 2nd positions are strongly constrained. About half of the 1st and 2nd codon positions are less variable than the most constrained 10% of 3rd positions (gray line).

When split by open reading frame (see Supp. Fig. 3 and Supp. Fig. 6), the most constrained regions are *ORF1b* and *M*, while *ORF3a, ORF6, ORF7a, ORF7b*, and *ORF8* show little evidence of constraint, consistent with frequently observed premature stop-mutations in some of these genes. *N* shows an intermediate pattern, possibly reflecting its mix of structured and unstructured regions. Variation at 3rd positions is common and comparable between genes. Only *E* shows slightly less variation at 3rd positions than other genes with a notable dip in the middle of the gene around codon 35 (see Fig. 6). Systematic difference in purifying selection on the amino acid sequence of different open reading frames have also been reported by **?**.

**FIG. 6.**
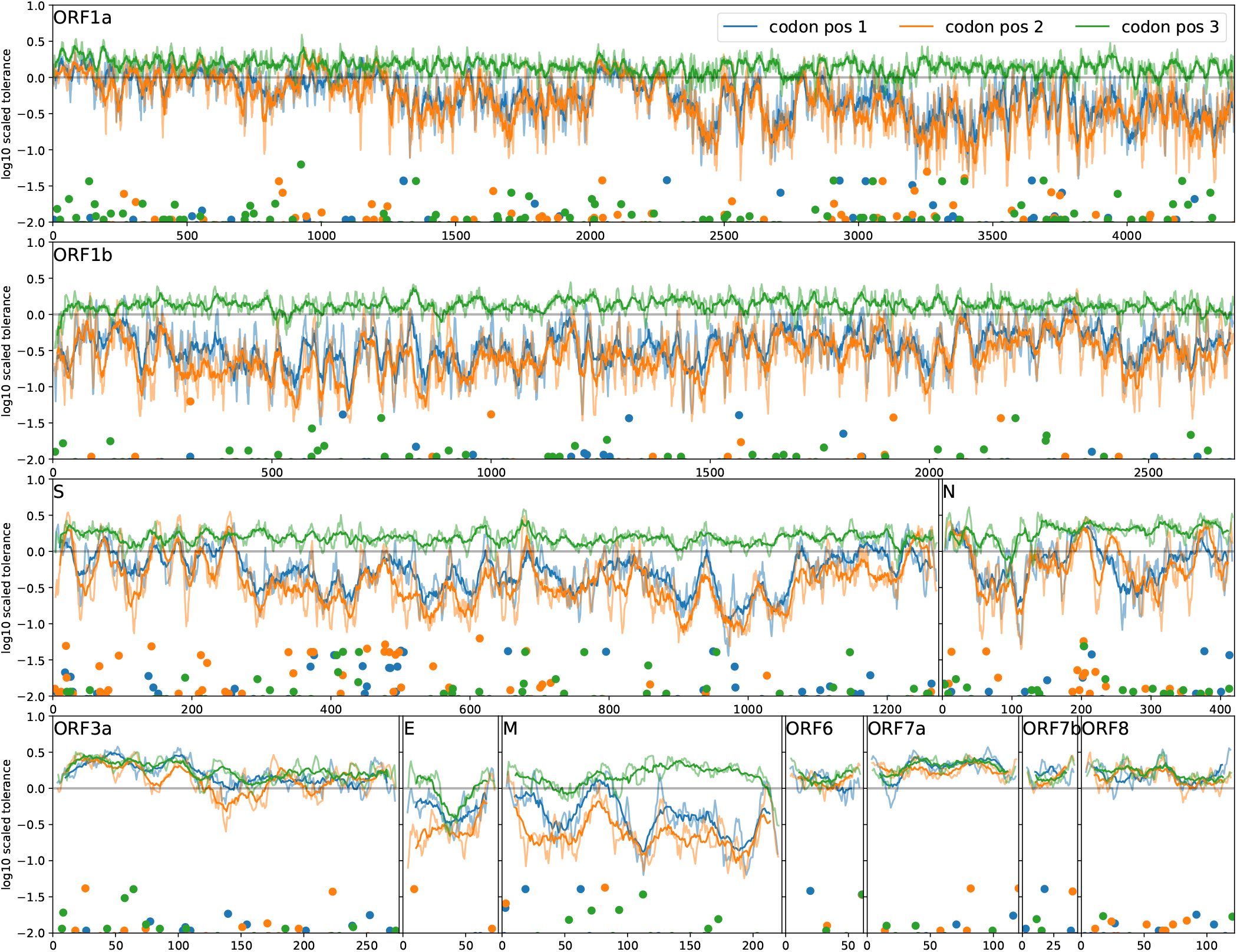
Landscape of selective constraint along the SARS-CoV-2 genome. Solid lines show sliding window smoothing of estimated mutational tolerance at 1st, 2nd, and 3rd positions for a window size of 20 and 7 (faint lines) sites. The markers at the bottom of the panels show qualitatively how many Pango lineages are differ at this position from the Wuhan-Hu-1 reference sequence.

## Discussion

The inferred evolutionary rate of RNA viruses often decreases with the time scale across which it is estimated (Ghafari *et al*., 2021; Wertheim and Kosakovsky Pond, 2011). This effect can be particularly pronounced at the beginning of an outbreak following a host switch and has been attributed to incomplete purifying selection or methodological issues leading to inflated measures of diversity (Ghafari *et al*., 2022; Meyer *et al*., 2015). In addition to segregating deleterious mutations, early viral evolution after a host switch can also be driven by anomalously fast adaptation. A dramatic change in environment, e.g. a host switch, likely results in many opportunities for mutations that increase fitness. Such transient increases in the rate of adaptation are common in experimental evolution (Elena and Sanjuán, 2007).

Here, I showed that the deceleration during the first six months of the pandemic is only observed for non-synonymous mutations. Since I analyzed evolution within short-lived clades of SARS-CoV-2 over a span of up to six months, purifying selection has comparable efficiency and should affect our estimates equally for all clades. Similarly, sequencing artifacts are expected to increase divergence at all time points and affect synonymous and non-synonymous rate in similar ways. The same applies to potential changes in the fidelity of the polymerase. Nevertheless, the estimated non-synonymous evolutionary rate of clades circulating in late 2019 and early 2020 is about twice as high as that of clades in 2021 and 2022, while the synonymous rate does not change (see Fig. 3 and Table I). One possible explanation is that the early evolutionary rate of SARS-CoV-2 was inflated by adaptive evolution.

Most of this apparent initial acceleration of within-variant evolution ceased in mid-2020 and even early variants like 20E accumulated non-synonymous changes at a rate of about 6 instead of 12 changes per year. By that time, the number of non-synonymous differences relative to the root of the tree was small (5 in the case of 20E) and it is implausible that this small change would have exhausted the pool of beneficial mutations. So why would the rate of adaptive evolution slow down? Maybe it is just chance, after all it is only clades 19A and 19B where the rate is substantially higher. Another possible explanation could be *diminishing returns* epistasis. Viruses with the S:D614G mutation showed faster replication kinetics compared to earlier variants (Korber *et al*., 2020). The virus might therefore be operating closer to the maximal capacity at which cells can produce virions, reducing the scope for further optimization. Other mutations then have smaller benefits, rise in frequency more slowly, and the effect of adaptation does not manifest itself over the life time of the clades studied here. Such diminishing returns epistasis has been observed in experimental evolution with yeast (Kryazhimskiy *et al*., 2014). Global epistasis of this nature can exist along side epistasis of specific amino acid changes as for example suggested for the S:N501Y mutation (Martin *et al*., 2021; **?**; **?**). Differences in diversification of SARS-CoV-2 with and without the S:D614G mutation where also observed in evolution experiments by Amicone *et al*. (2022), who concluded that these differences were not due to a change in base line mutation rate but had selective origin.

An evolutionary rate of 6 synonymous changes per year at around 9700 positions corresponds to a per-site evolutionary rate of 6.2 × 10^−4^/site/year or 1.7 ×10^−6^/day, slightly higher than the estimated base-line mutation rate of 1.3 × 10^−6^/day (Amicone *et al*., 2022). The total evolutionary rate within variants after mid 2020 (starting with 20E) is in the range of 9-16 changes per year, corresponding to a per site rate of 3 − 5 ×10^−4^/site/year, consistent with recent estimates by Hill *et al*. (2022) and Tay *et al*. (2022).

When considering only the putative founder genotype and date of origin of each variant, all variants so far are compatible with a back-bone evolutionary rate of 32 changes per year, corresponding to an per site rate of around 10^−3^/site/year (see Fig. 3). This rate is a composite of episode of cryptic accelerated evolution (possibly in chronically infected individuals), and regular transmission chains of acute infections. This estimate for the average rate is thus not inconsistent with results by Hill *et al*. (2022) and Tay *et al*. (2022), who estimated even higher rates specifically for the branches that gave rise to variants of concern. Different clades and variants probably emerged in different ways under different circumstances. Nevertheless, all clades and variants are compatible with a single “back-bone” molecular clock that runs substantially faster than the “within variant clock”. This accelerated back-bone clock is likely driven by exponential amplification of beneficial mutations.

In addition to neutral and potentially adaptive mutations, I also quantified purifying selection on the SARS-CoV-2 genome. By analyzing the rate of mutations that spread only on short times scales within fine-grained Pango lineages, I estimated the level of constraint on different parts of the SARS-CoV-genome. The great majority of 3rd positions in codons – at which most mutations are synonymous – do not show strong signatures of conservation. The major open reading frames *ORF1ab, S, N, E* and *M* show clear signatures of purifying selection with around 50% of sites being more constrained than the most constrained 10% of 3rd positions. Purifying selection analyzed here is operating over the time during which rare lineage circulate, which is typically a few weeks. On shorter time scales, more strongly deleterious mutations will be observed. On longer time scales, weaker deleterious effects can be quantified. The degree of purifying selection is consistent with a similar analysis in HIV-1 that also inferred small fitness costs for most synonymous mutations, while half of the non-synonymous mutations are so deleterious that they are not even observed on short timescales (Zanini *et al*., 2017).

The SARS-CoV-2 genes *ORF3a, ORF6, ORF7a, ORF7b* and *ORF8* show little global signal of constraint and 1st and 2nd positions are as variable as 3rd positions (see Fig. 5). This lack of strong purifying selection in these ORFs does not necessarily imply that they don’t matter or are not possible loci of adaptation – but the exact amino acid sequence does not seem essential for their function. Only a few regions show a clear signal of conservation at 3rd positions, notably a central region of *E* and the ribosome slippage region at the beginning of *ORF1b*.

The heterogeneity in evolutionary rates and the combination of adaptive evolution, approximately neutral mutations, and purifying selection complicate the interpretation of phylodynamic analysis. Phylodynamics typically assumes that the mutation process is independent of the spread and epidemiology and that different sites evolve independently. These assumptions are (approximately) true for neutral mutations that occur along every lineage with the same rate. Strongly deleterious mutations don’t spread and are only observed on terminal branches, similar to sequencing errors. Weakly deleterious mutations can spread, but lineages that carry them tend to die out. Overall, the deleterious mutations result in increased diversity on short time scales compared to longer ones, which can lead to time-dependent effective evolutionary rates (Wertheim and Kosakovsky Pond, 2011).

The effects of adaptive evolution are harder to control and account for. Since the number of sites that allow beneficial mutations is small, adaptive evolution tends to be very stochastic – it is not the typical events, but the rare and extreme events that determine the course of adaptive evolution. Unlike neutral evolution, the rate of adaptive evolution depends on the population size, stochastic nature of the transmission process, the environment, and previous adaptation (Neher, 2013). These factors make extrapolation – whether it is reconstruction of past events or scenarios of future evolution – uncertain. Standard phylogenetic or phylodynamic methods will underestimate this uncertainty.

If the excess of non-synonymous changes on the pandemic scale compared to within clade evolution is due to adaptive evolution, it would imply an adaptive rate of about 20 changes per year (plus insertion/deletions not considered here). This process is likely to continue even in the more and more diverse immunological landscape of the human population in which the viral population spread and adapts. But it is unclear whether SARS-CoV-2 continues to evolve in a saltatory fashion with repeated emergence of highly mutated variants, or whether we will see a transition to a more gradual adaptive process. We currently observe multiple lineages with several amino acid changes and moderate transmission advantages emerging in a more stepwise fashion within 21L, 22A, 22B (BA.2, BA.4, BA.5), which could indicated a shift to a more influenza-like evolution.

## Materials and Methods

### Data

This analysis is based on SARS-CoV-2 sequencing data shared by researchers around the world via GISAID (Shu and McCauley, 2017) (EPISET_ID *EPI_SET_220812zf*). The bulk of the analysis can be reproduced using SARS-CoV-2 genomes shared via INSDC. These data are available from Nextstrain at http://data.nextstrain.org/files/ncov/open/metadata.tsv.gz and http://data.nextstrain.org/files/ncov/open/nextclade.tsv.gz. However, for some clades and variants, coverage in INSDC is low.

### Analysis

The code used to run this analysis is publicly available at https://github.com/neherlab/SC2_variant_rates. It is organized as a snakemake workflow (Köster and Rahmann, 2012). I used Nextclade data *2022-07-26T12:00:00Z* with reference sequence MN908947 (Wuhan-Hu-1). The codon analysis ignores *ORF9b* which is fully contained in *N* but in a different reading frame. Tabular files with estimates of evolutionary rates, the mutation distribution, and fitness costs are included in the git repository.

The fitness costs of each positions are estimated from the Nextclade columns *unlabeledSubstitutions* and *reversionSubstitutions* for each sequence that has QC-status good. For each Pango lineage, I count how often each specific mutation was observed. I then calculate how many lineages show minor variation for each mutation. To avoid loss of signal through sequencing errors, I drop lineages at which only a single mutation at this position is observed. To turn this count into a measure of mutational tolerance, I normalize the number of lineages with minor variation by the mutation rate away from the ancestral base at this position within the lineage.

## Acknowledgements

I gratefully acknowledge the work by 1000s of scientists and healthcare professionals that collect the samples, generate, and share the sequence data on which this analysis is built. I am also grateful to Jesse Bloom, Louis du Plessis, and Florence Débarre for valuable feed-back on the manuscript and to Cornelius Roemer and Emma Hodcroft for stimulating discussions on many aspects of SARS-CoV-2 evolution.

## Appendix: Supplementary figures

**Figure S 1.**
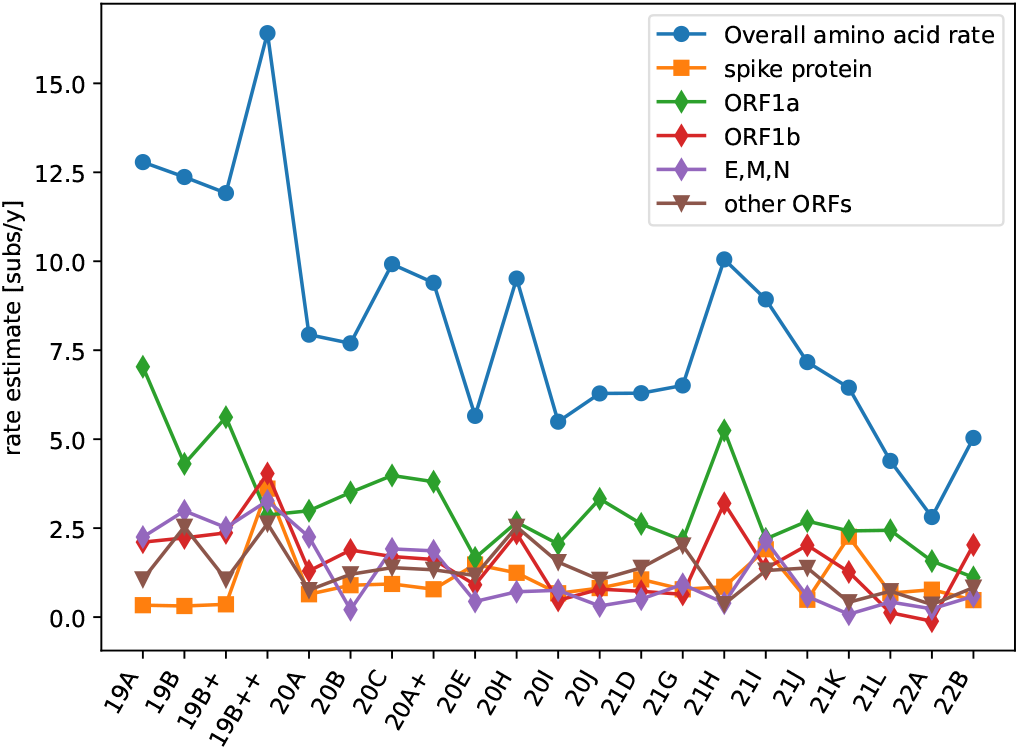
Within-clade non-synonymous evolution in different parts of the genome. The decreasing trend in the within-clade non-synonymous rate of evolution is most evident in the largest *ORF1ab*. Accessory proteins and *N,E,M* show a weak trend. Clades are ordered alphabetically, which corresponds to their order of designation, which in turn is very similar to their order of emergence.

**Figure S 2.**
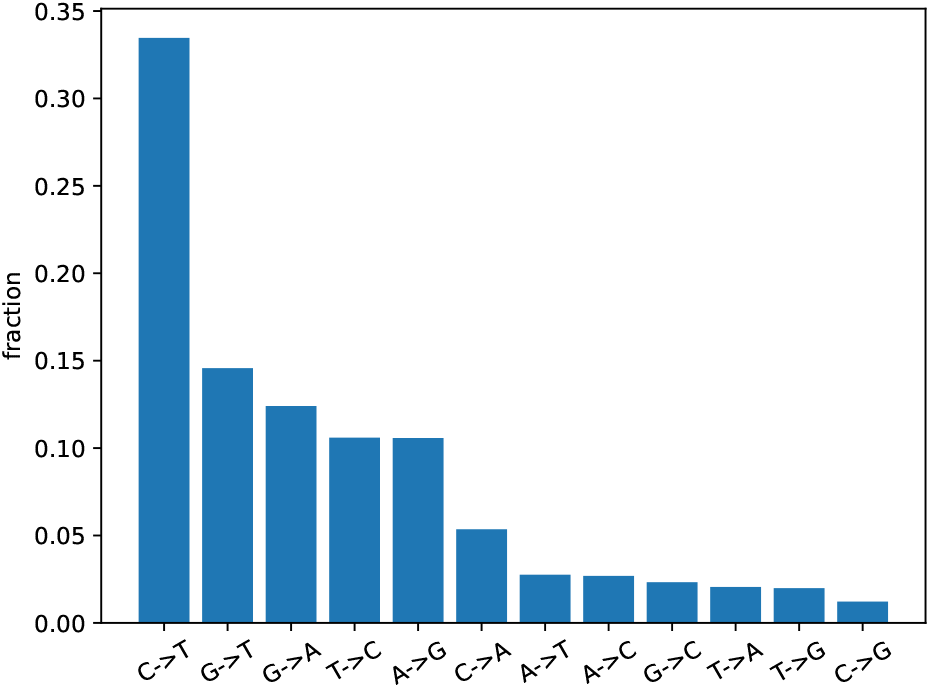
The relative rate of different mutations in SARS-CoV-2. These rates are measured from rare low frequency mutations probably subject to little purifying selection.

**Figure S 3.**
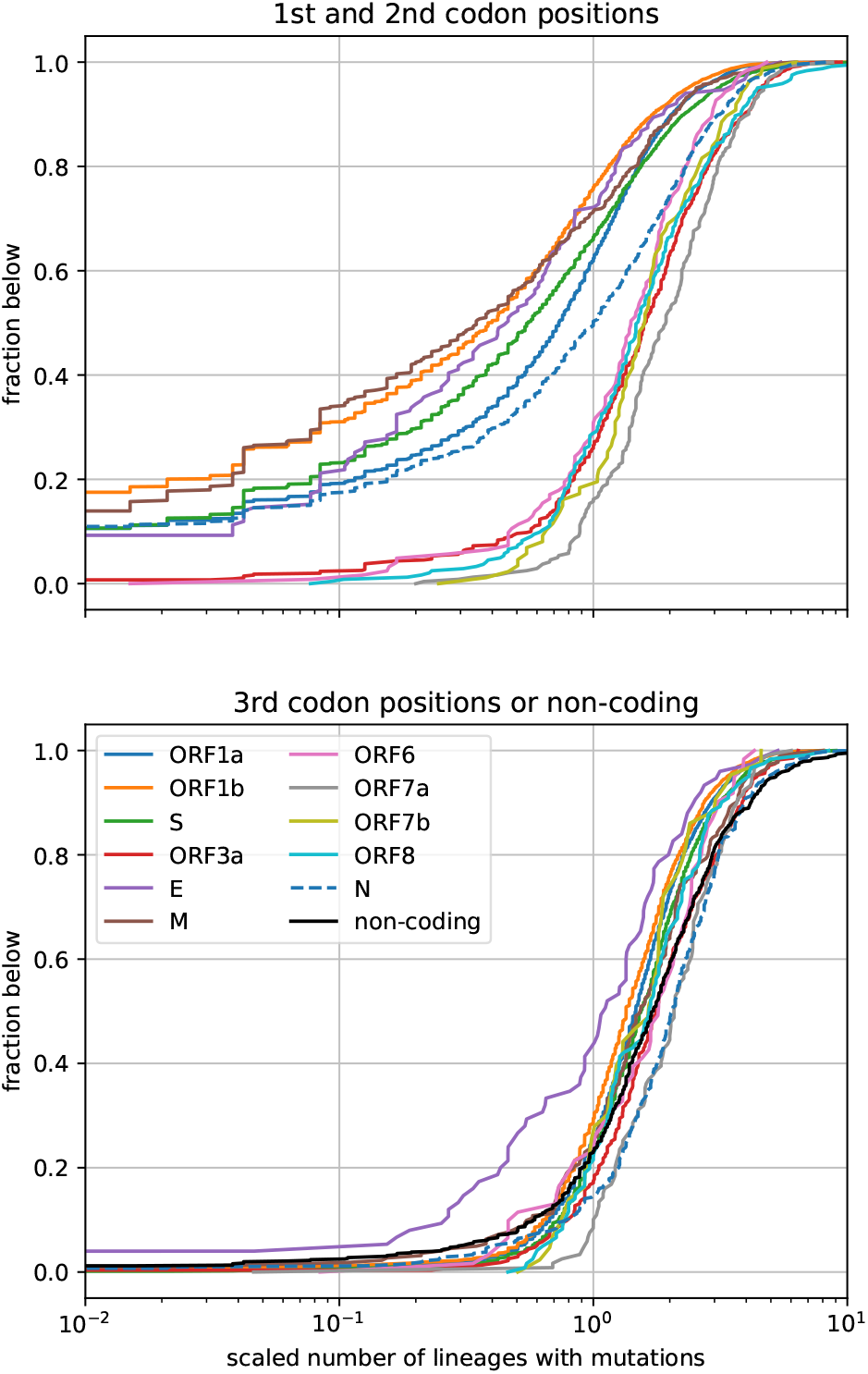
Constraints on SARS-CoV-2 mutations by gene. The top panel quantifies constraint at 1st and 2nd positions in codons of open reading frames. The bottom panel show the analogous distributions at 3rd codon positions. The latter distributions are very similar across genes, with only *E* showing somewhat less variation. In contrast, mutation tolerance at 1st and 2nd positions differs markedly between genes. In ORF3a, ORF6, ORF7a, ORF7b, and ORF8 the distribution of mutations at 1st and 2nd positions is very similar to the distribution at 3rd positions, while the remaining genes show clear signs of constraint.

## Appendix: Divergence summaries

**Figure S 4.**
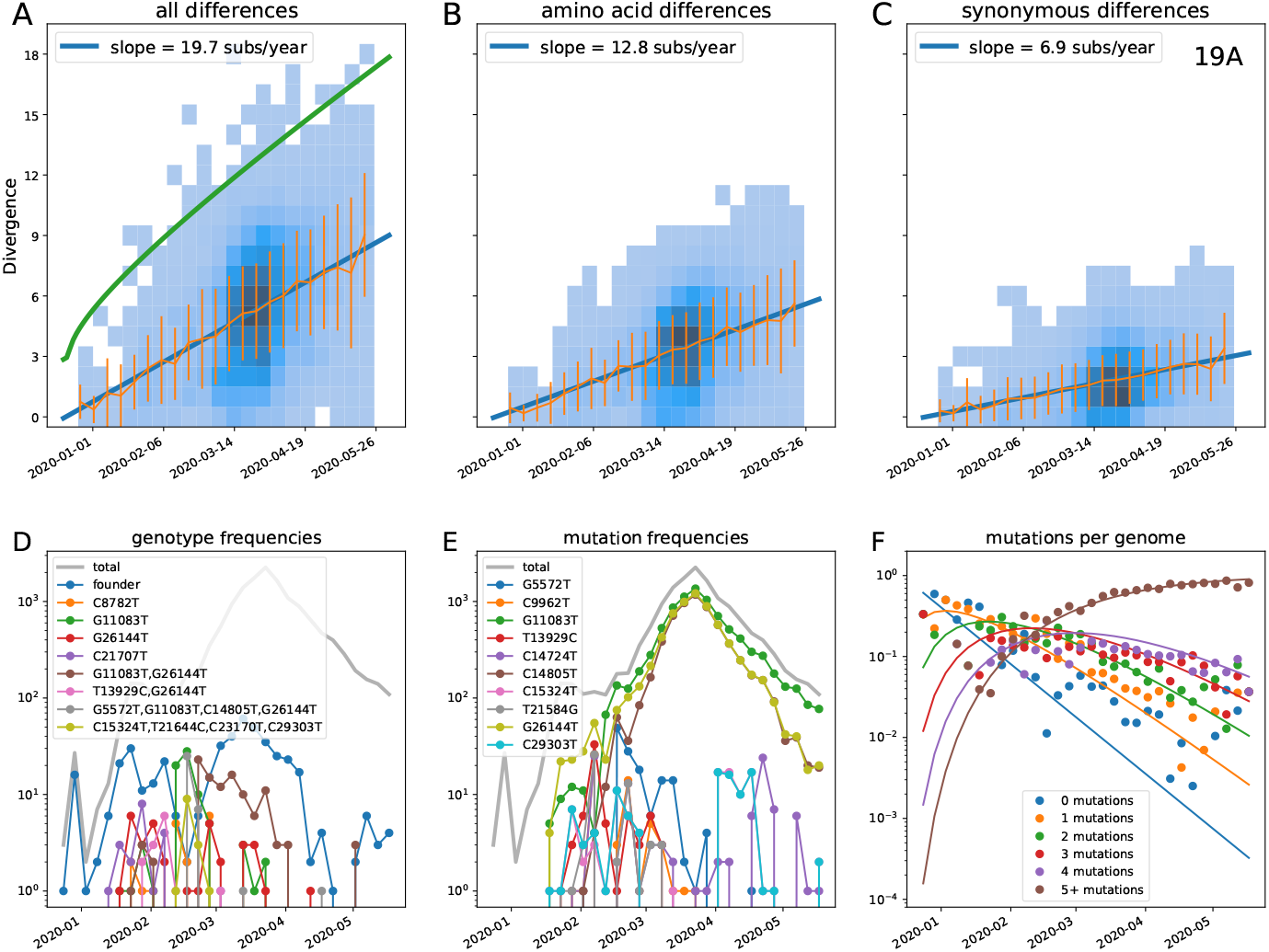
Divergence increases linearly with time in clade 19A. Top: Each panel shows the number of within-clade mutations (total (A), amino acid changing (B), synonymous (C)) as a function of time. The green line in panel A indicates the divergence cut-off, panels B&C only show sequences that pass the divergence filter. Each panel also shows mean *±* standard deviation and a weighted linear fit. Analogous figures for all clades considered are included in the appendix. Bottom: Panels D and E show the prevalence of specific genotypes (D) and specific mutations (E). In the case shown, the founder genotype initially dominates and no daughter genotype or mutation dominate. Panel F shows a Poisson model to the breakdown of the population into genotypes with different number of mutations over time.

**Figure S 5.**
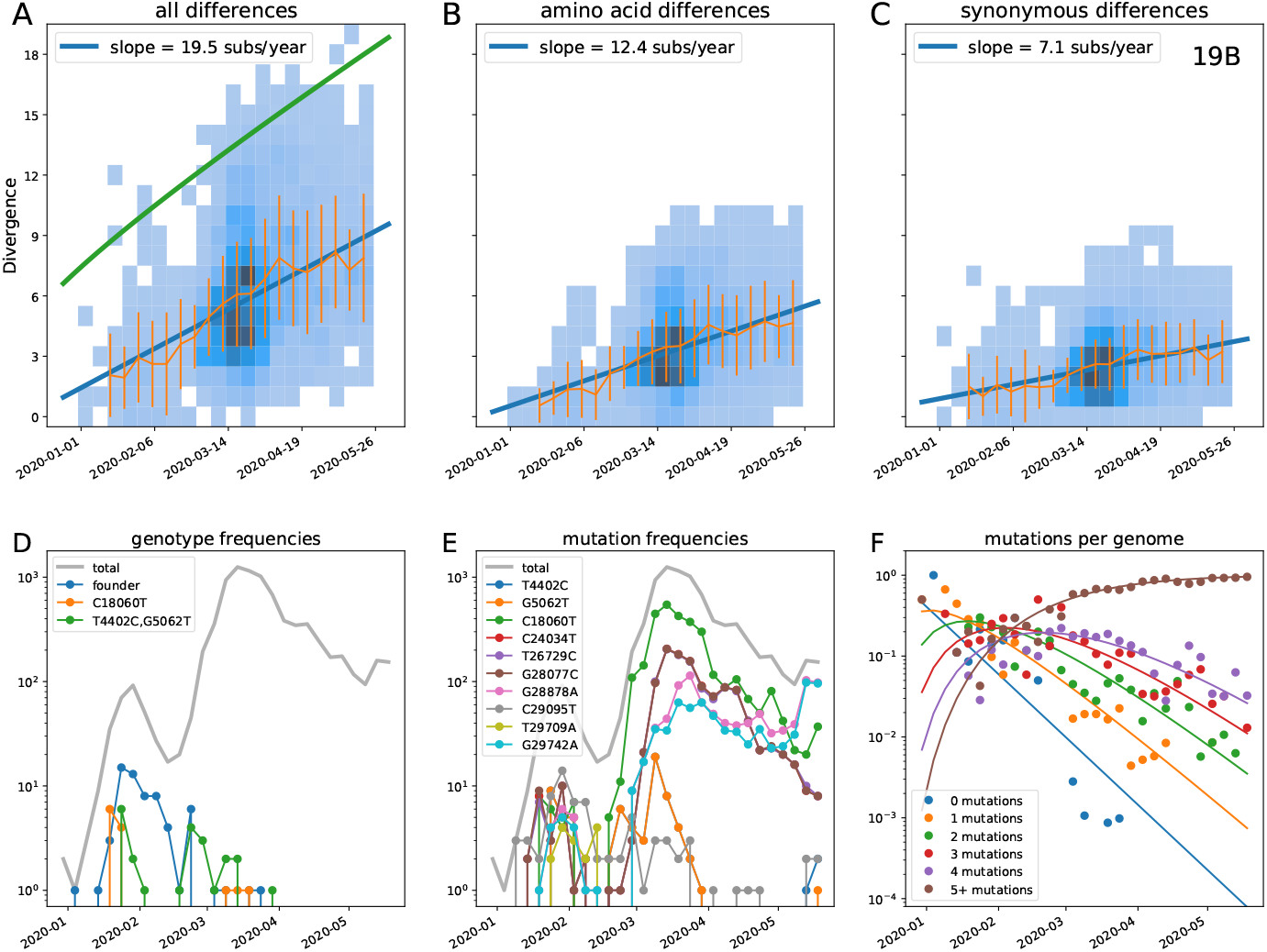
Divergence increases linearly with time in clade 19B.

**Figure S 6.**
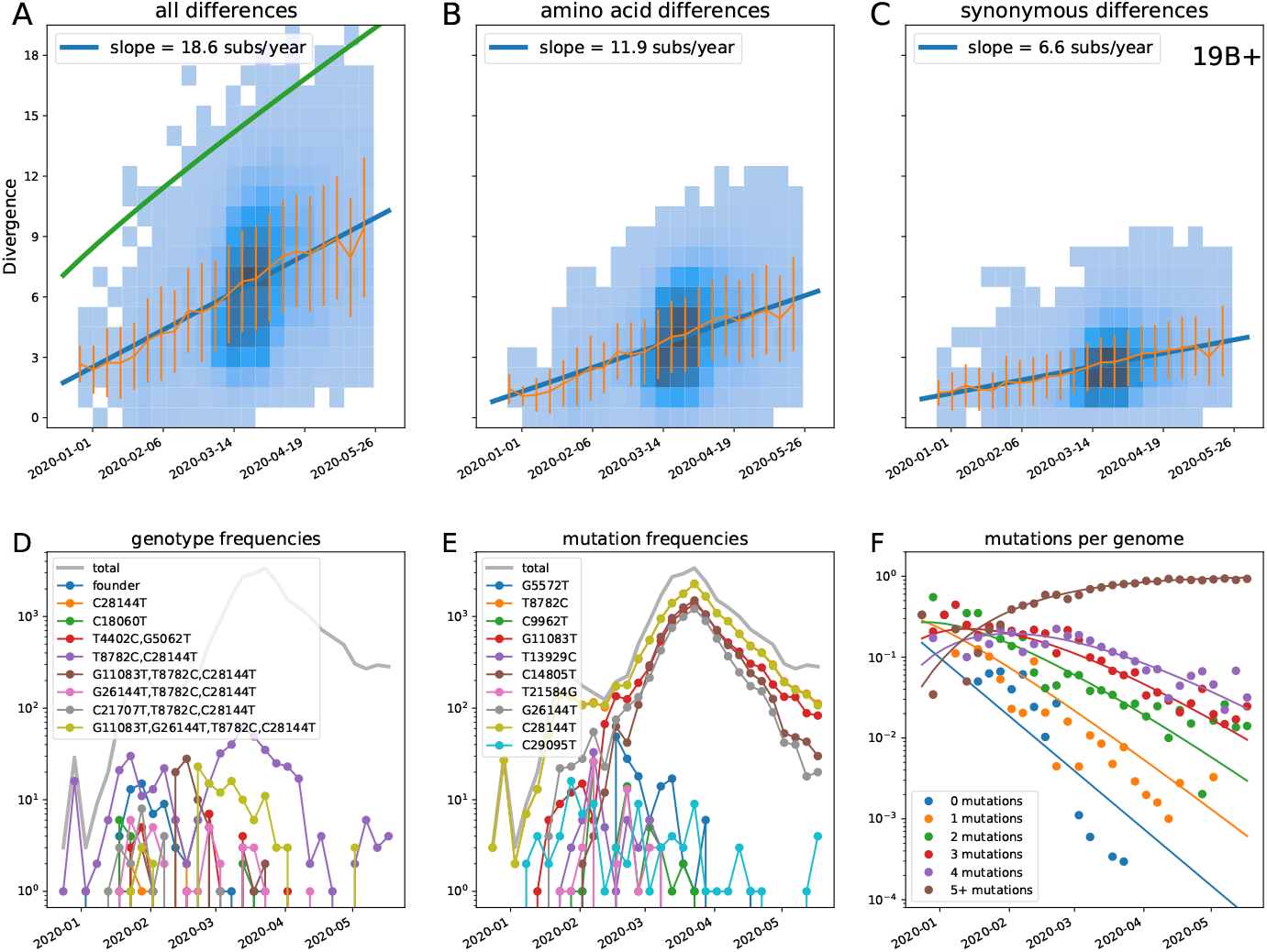
Divergence increases linearly with time in clade 19B+. This figure contains sequences in clades 19 A and B rooted on clade 19B.

**Figure S 7.**
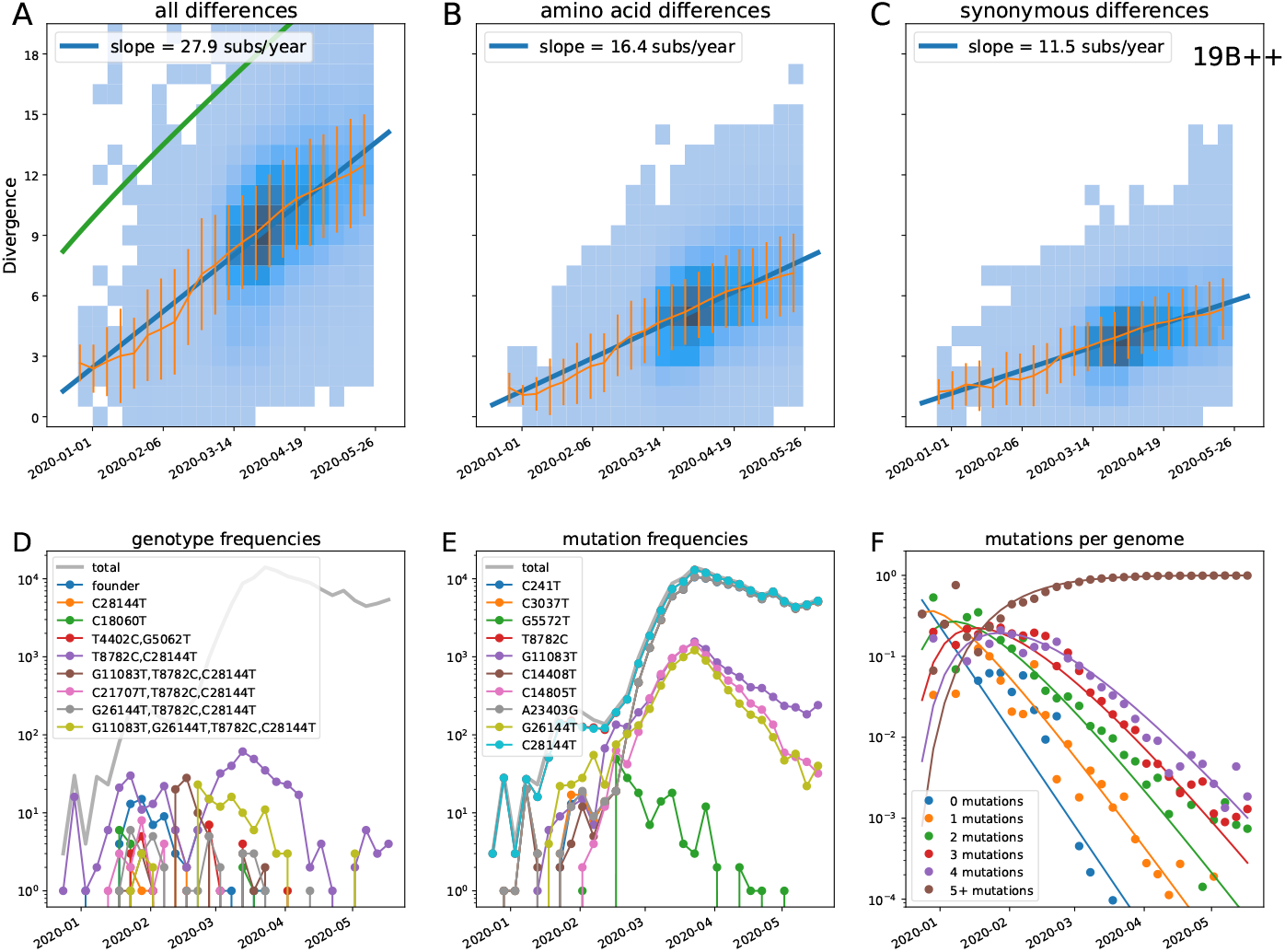
Divergence increases linearly with time in clade 19B++. This figure contains sequences in clades 19A, 19B, 20A, 20B, 20C, and 20D rooted on clade 19B.

**Figure S 8.**
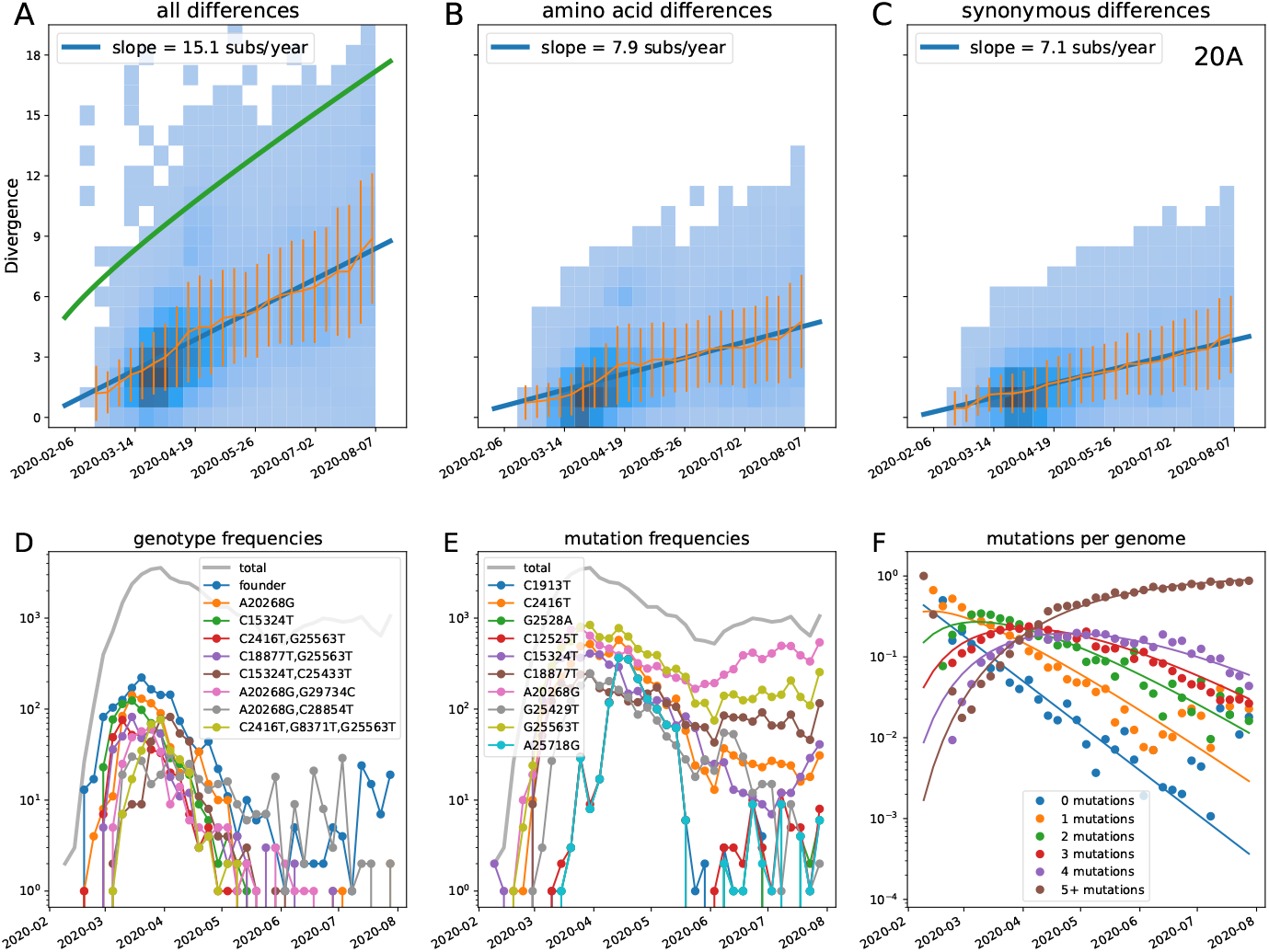
Divergence increases linearly with time in clade 20A.

**Figure S 9.**
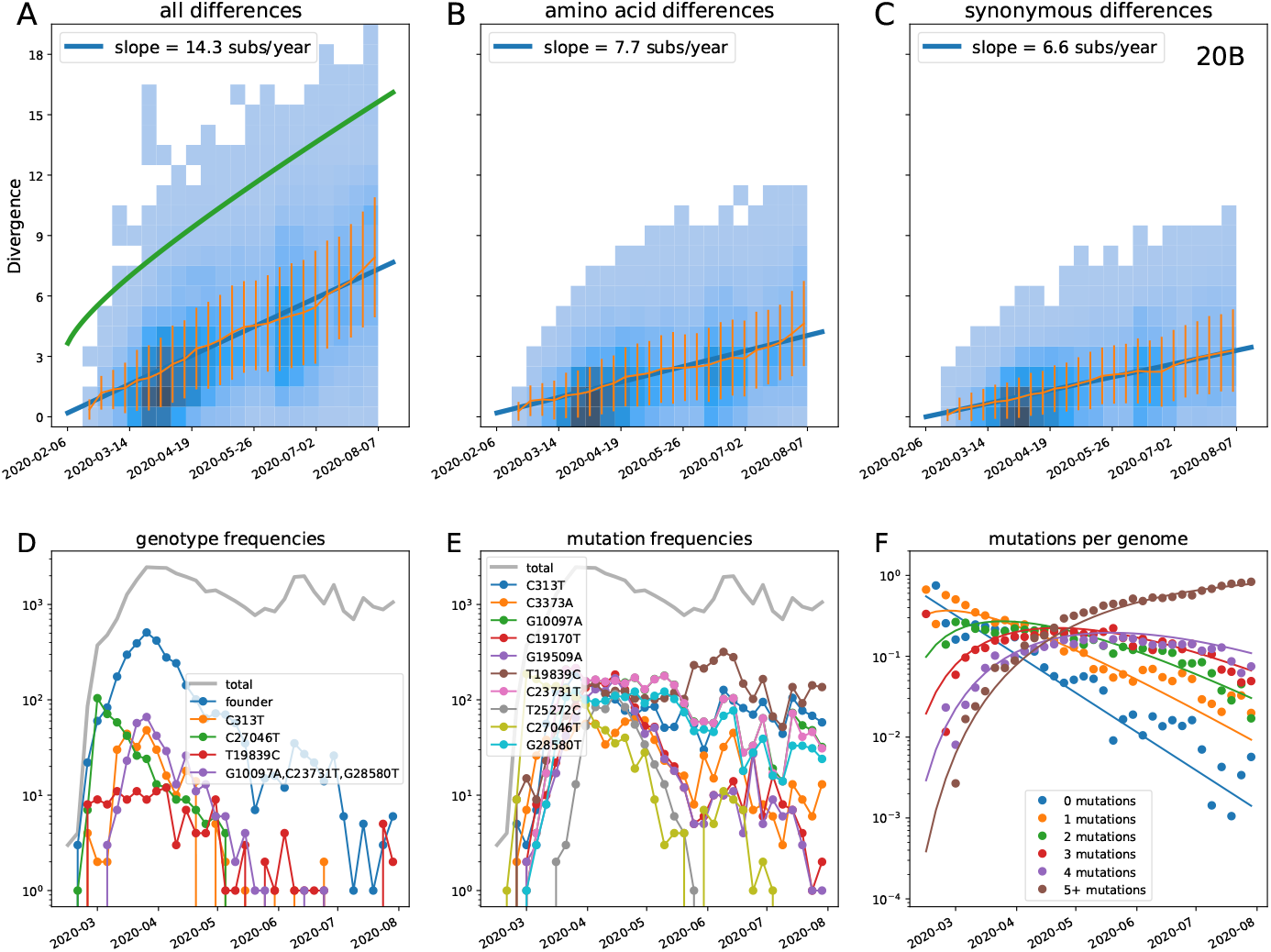
Divergence increases linearly with time in clade 20B.

**Figure S 10.**
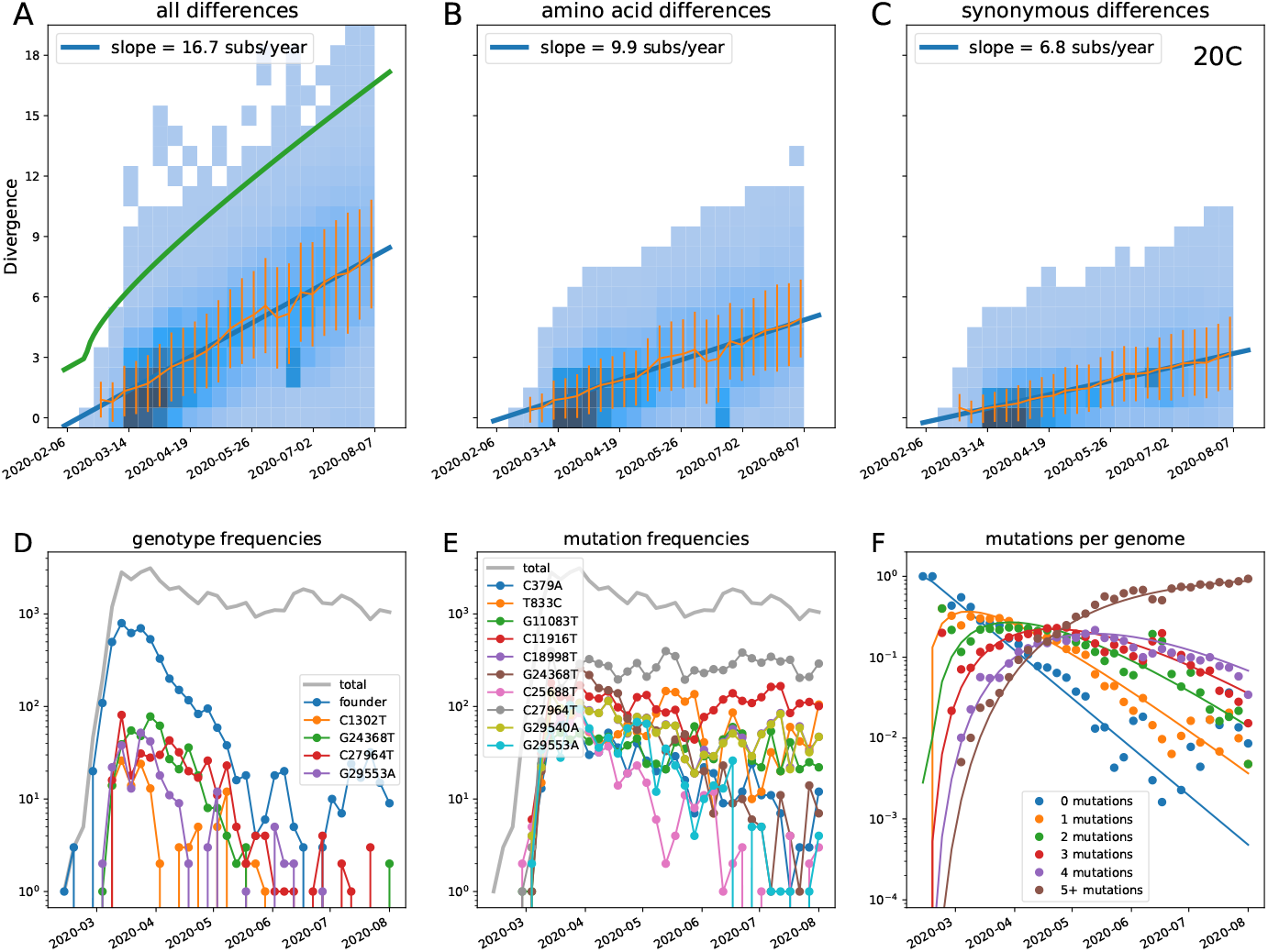
Divergence increases linearly with time in clade 20C.

**Figure S 11.**
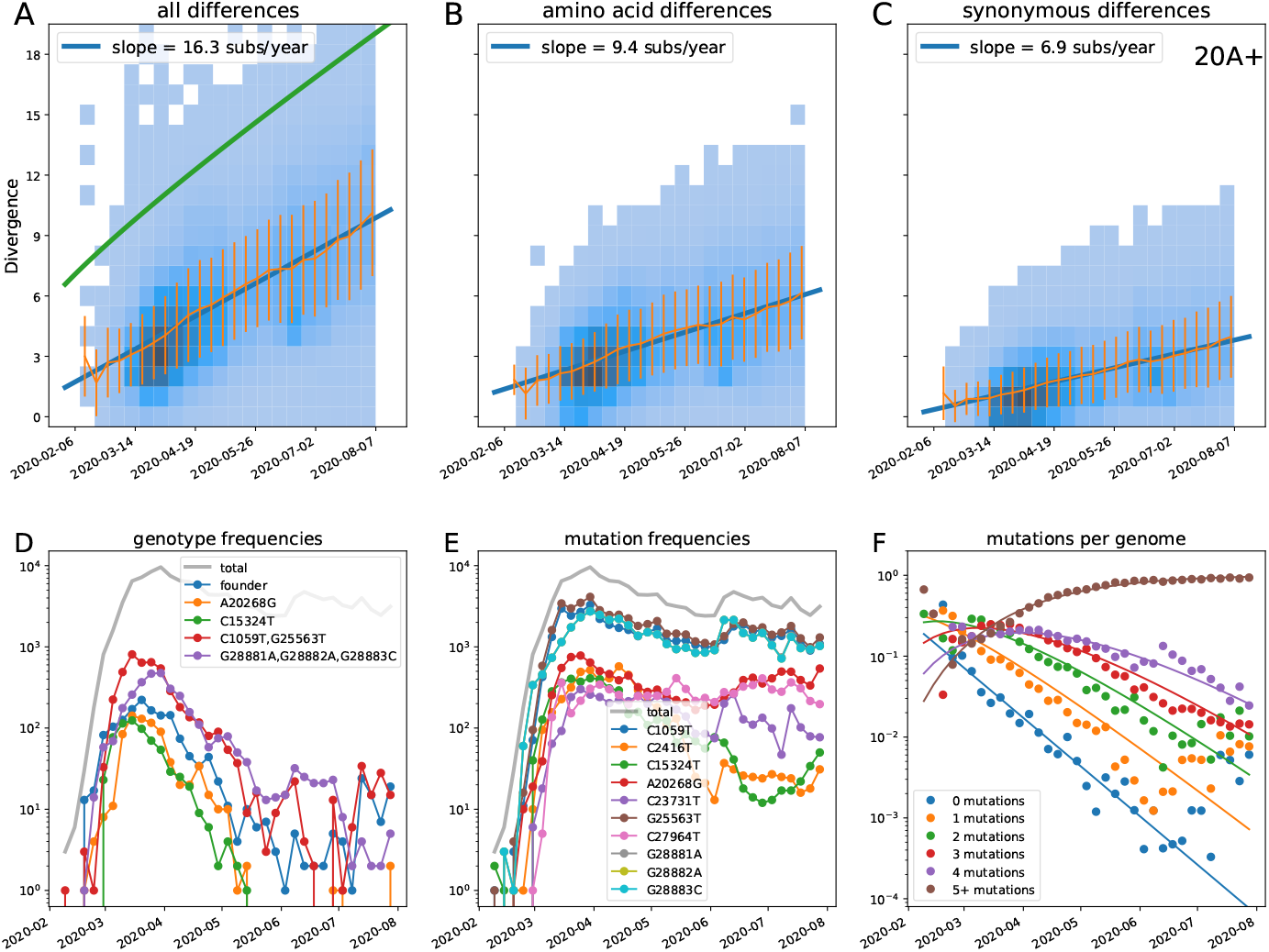
Divergence increases linearly with time in clade 20A+. This figure contains sequences in clades 20A,B,C,D rooted on clade 20A.

**Figure S 12.**
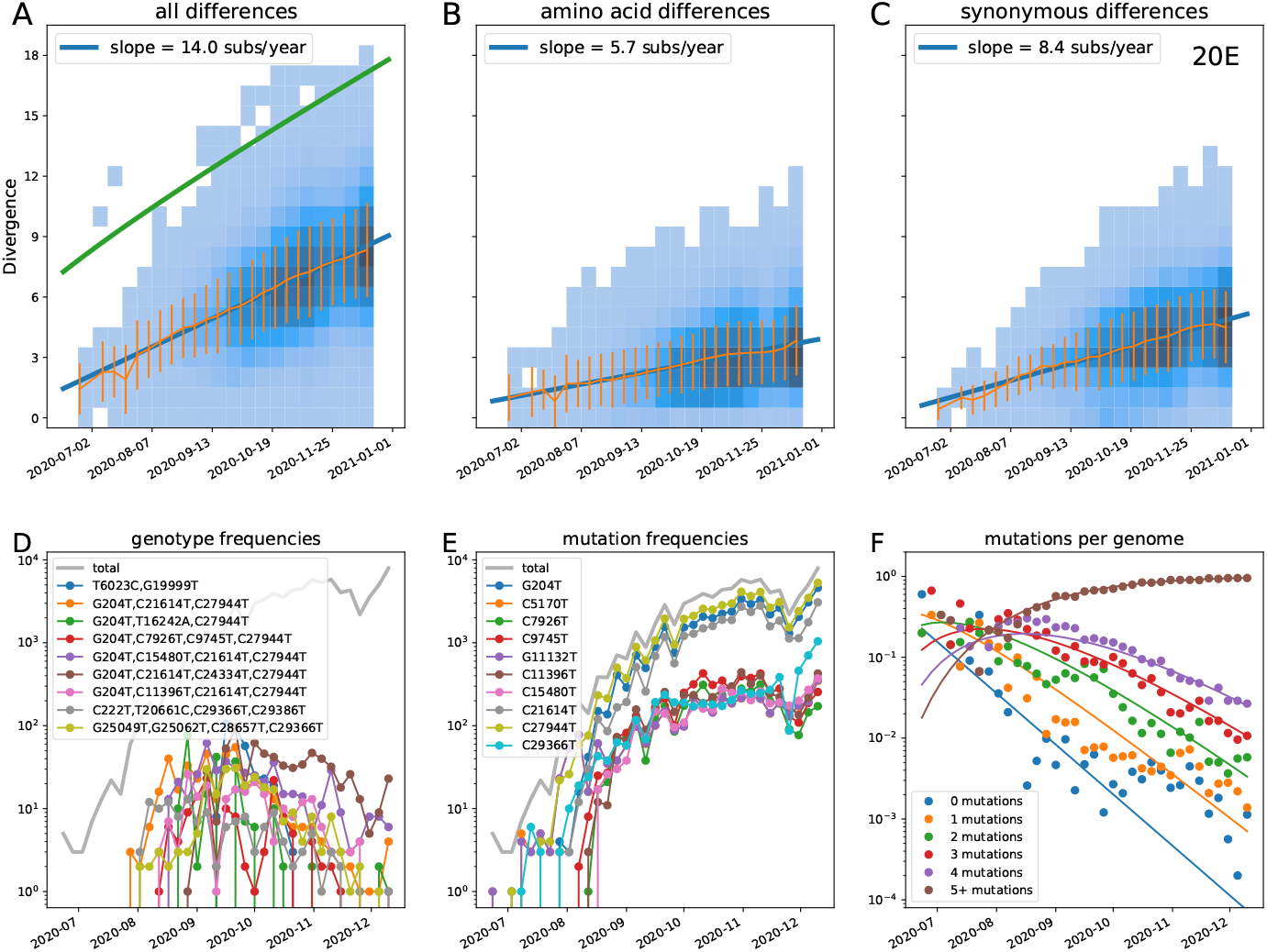
Divergence increases linearly with time in clade 20E.

**Figure S 13.**
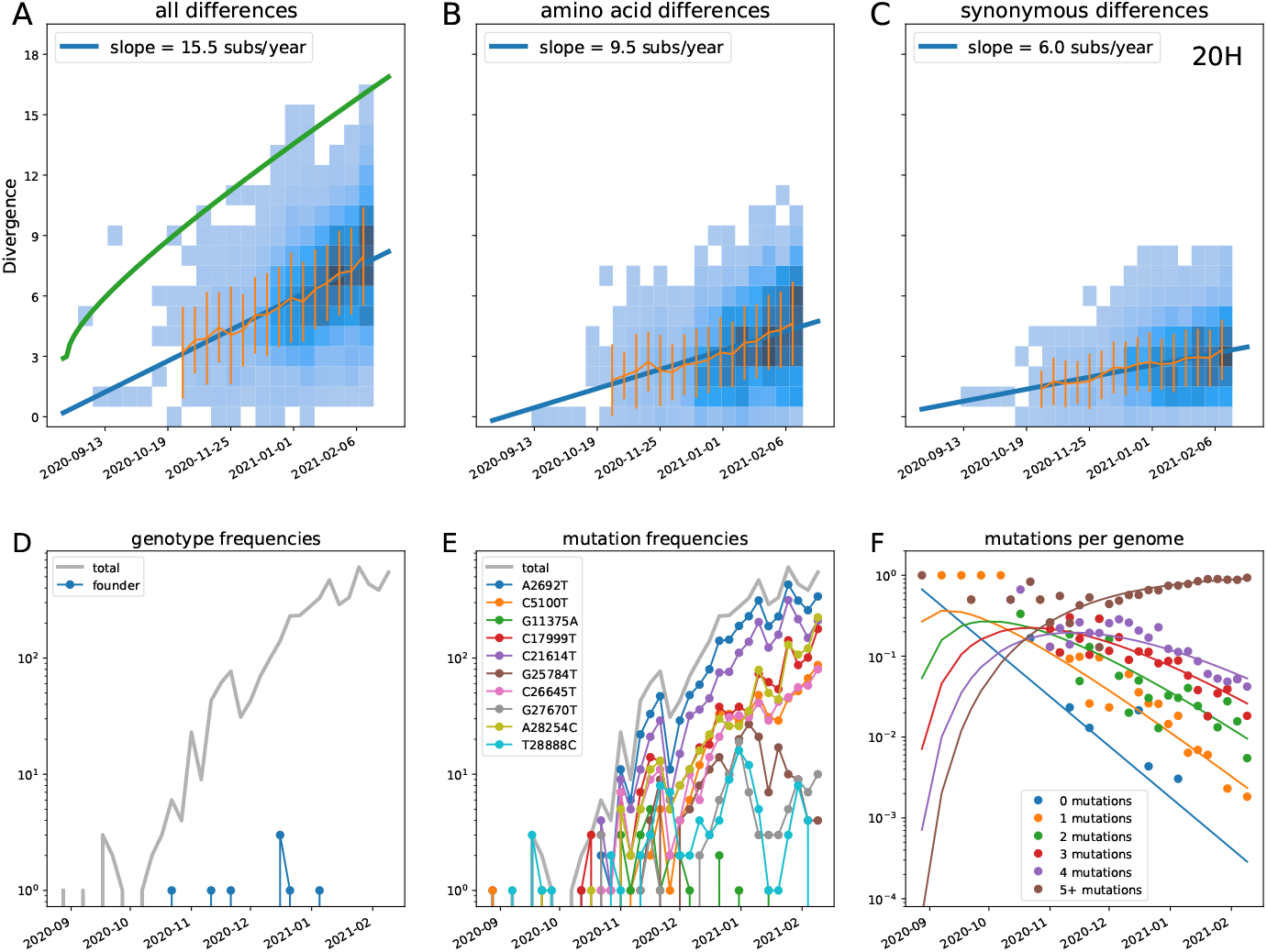
Divergence increases linearly with time in clade 20H (Beta).

**Figure S 14.**
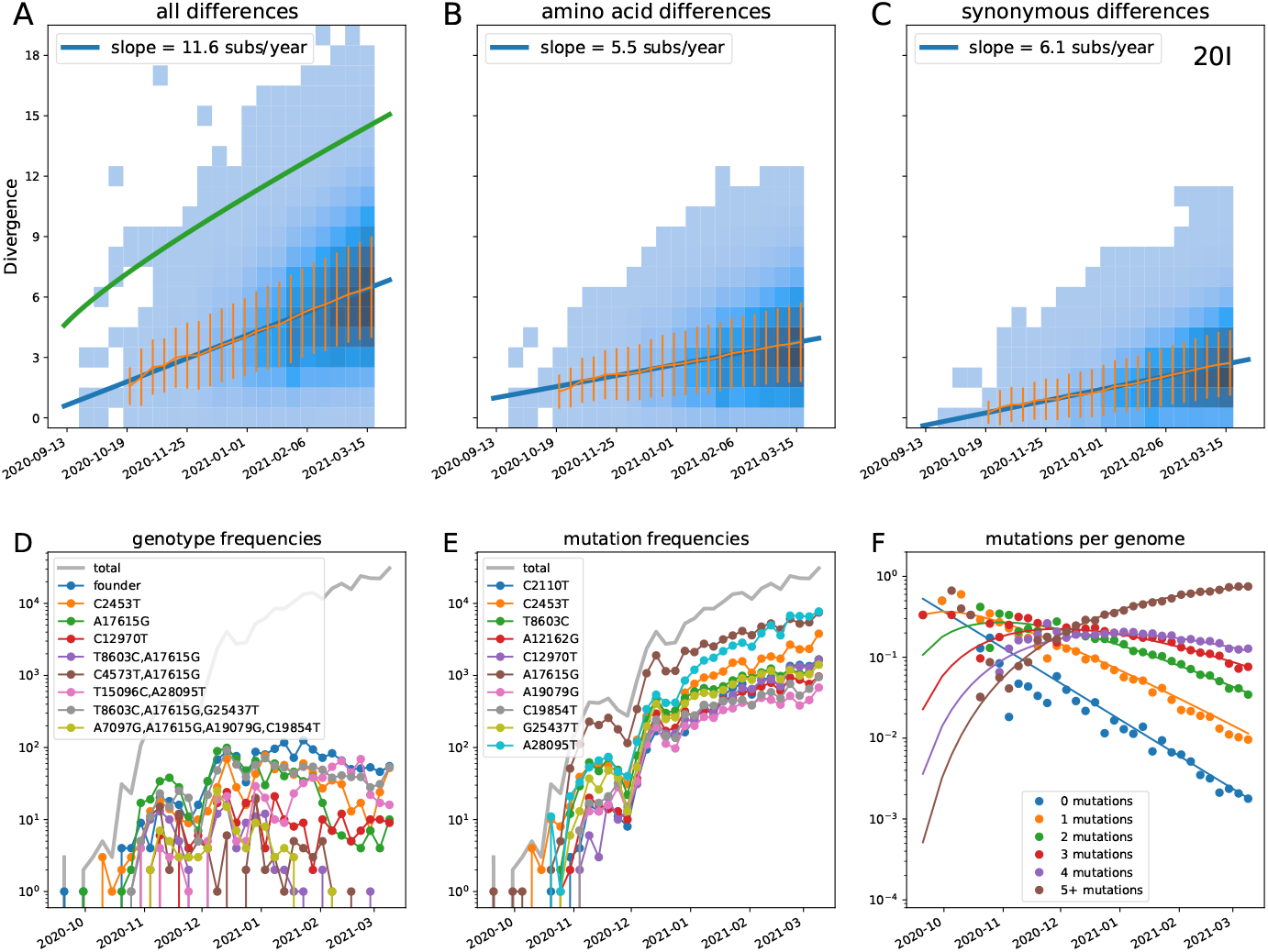
Divergence increases linearly with time in clade 20I (Alpha).

**Figure S 15.**
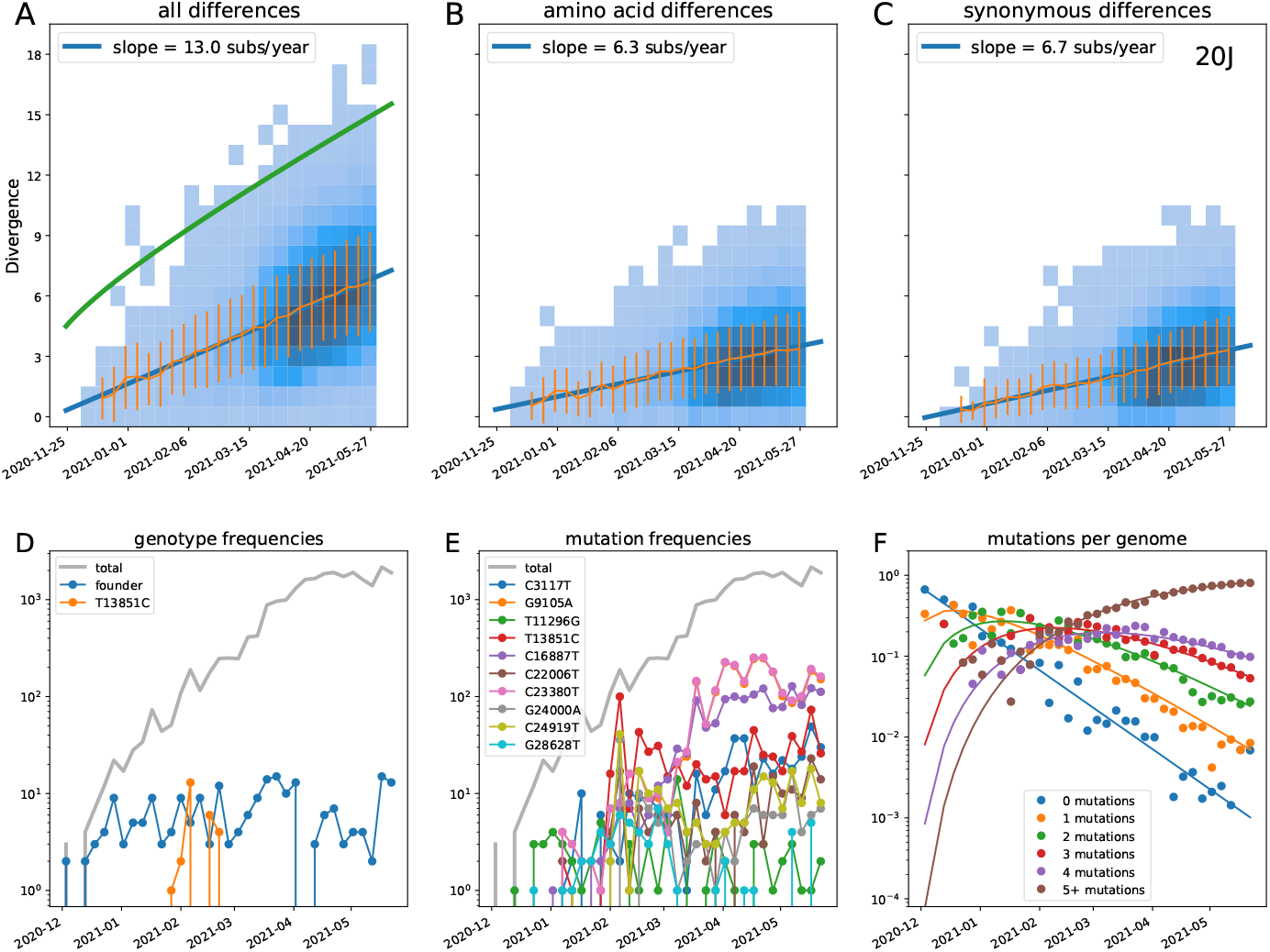
Divergence increases linearly with time in clade 20J (Gamma).

**Figure S 16.**
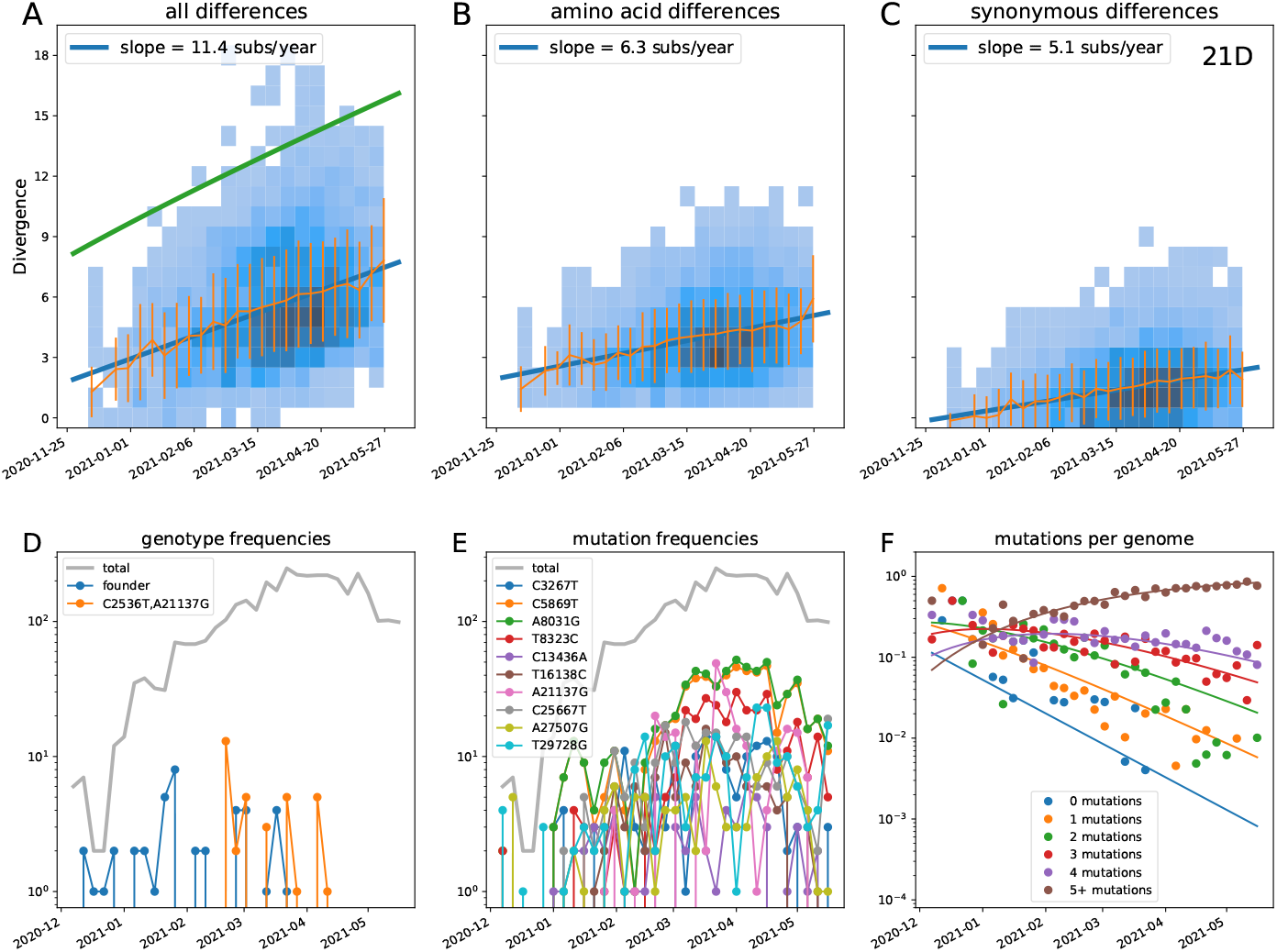
Divergence increases linearly with time in clade 21D (Eta).

**Figure S 17.**
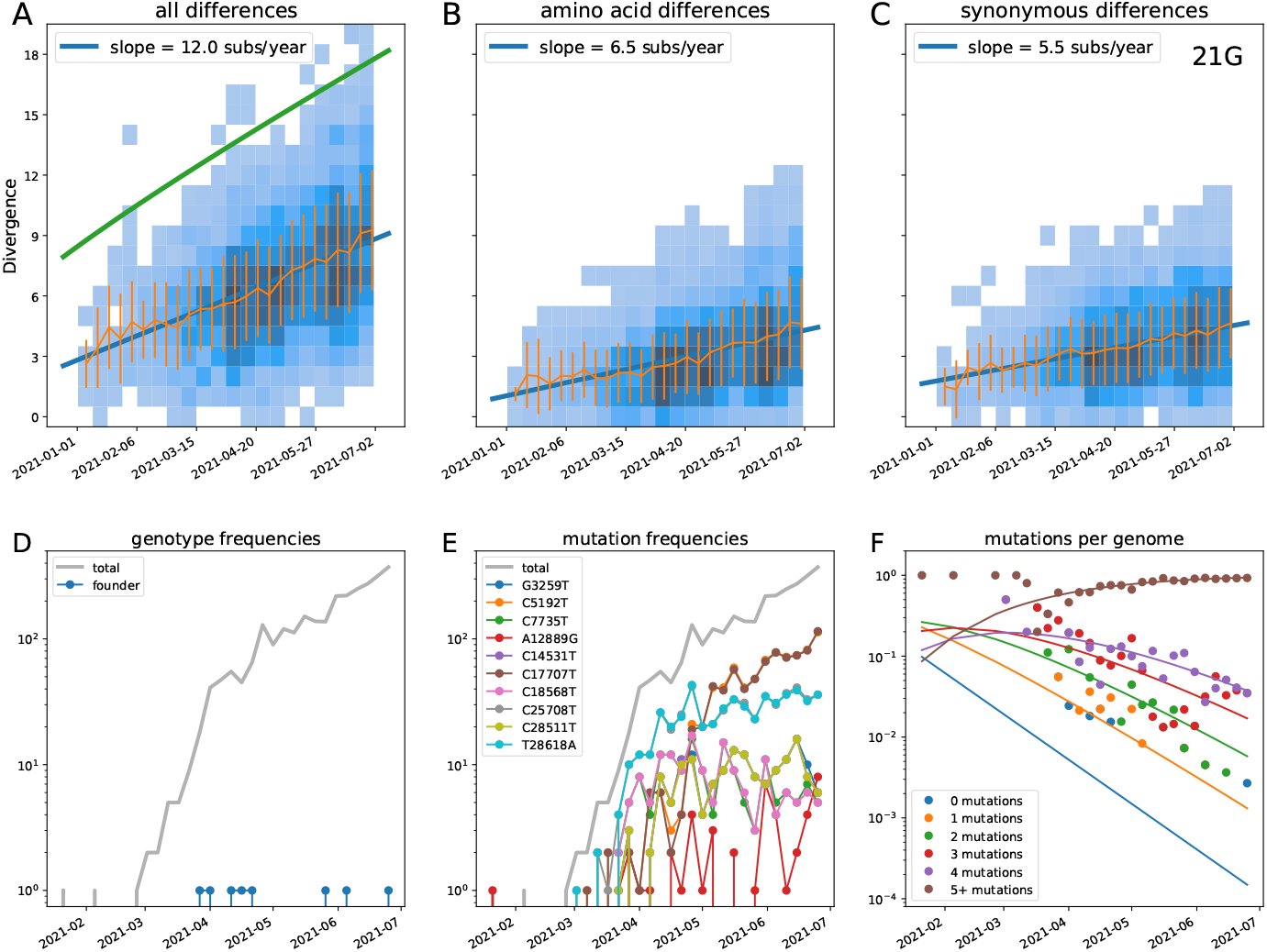
Divergence increases linearly with time in clade 21G (Lambda).

**Figure S 18.**
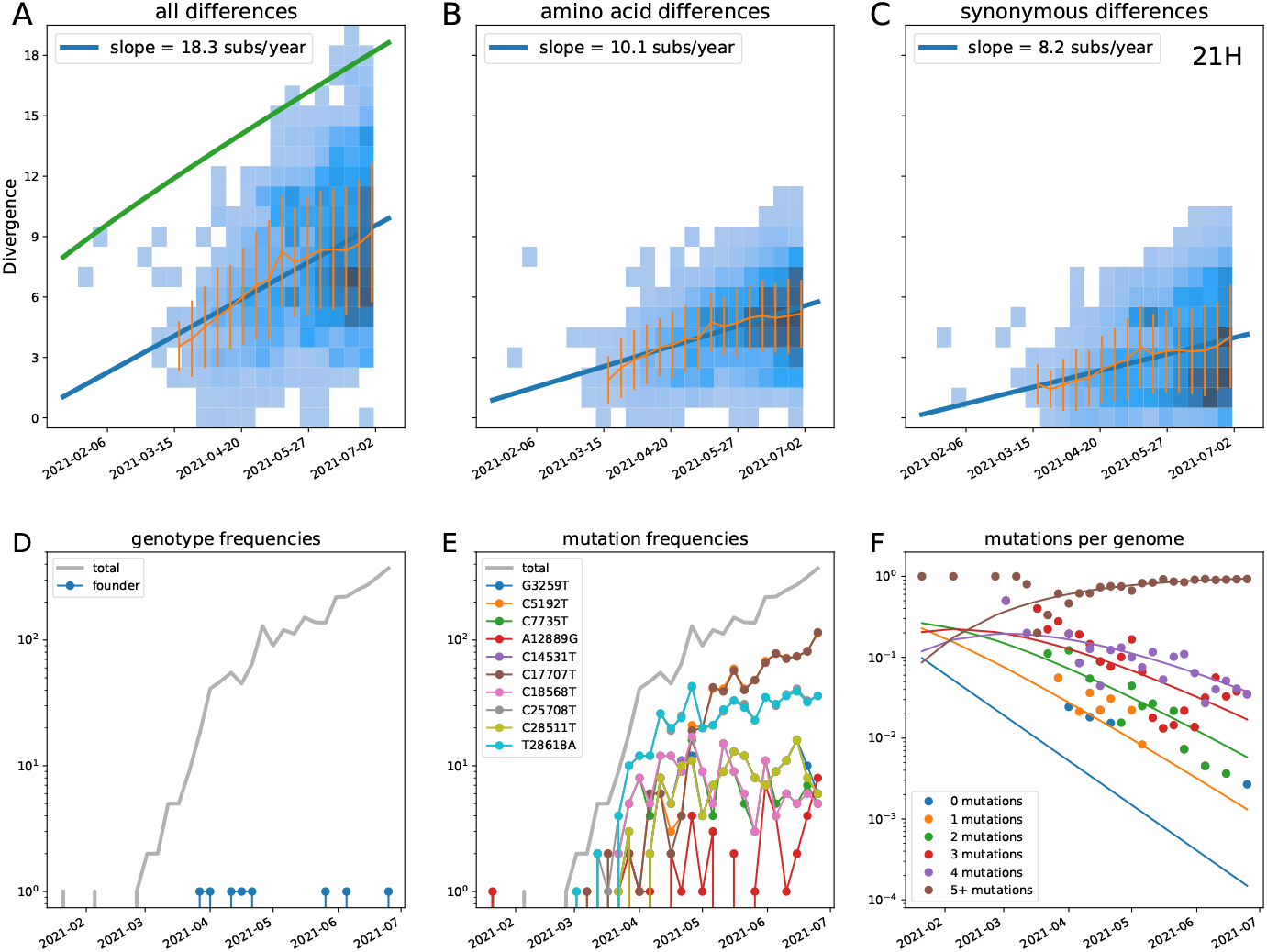
Divergence increases linearly with time in clade 21H (Mu).

**Figure S 19.**
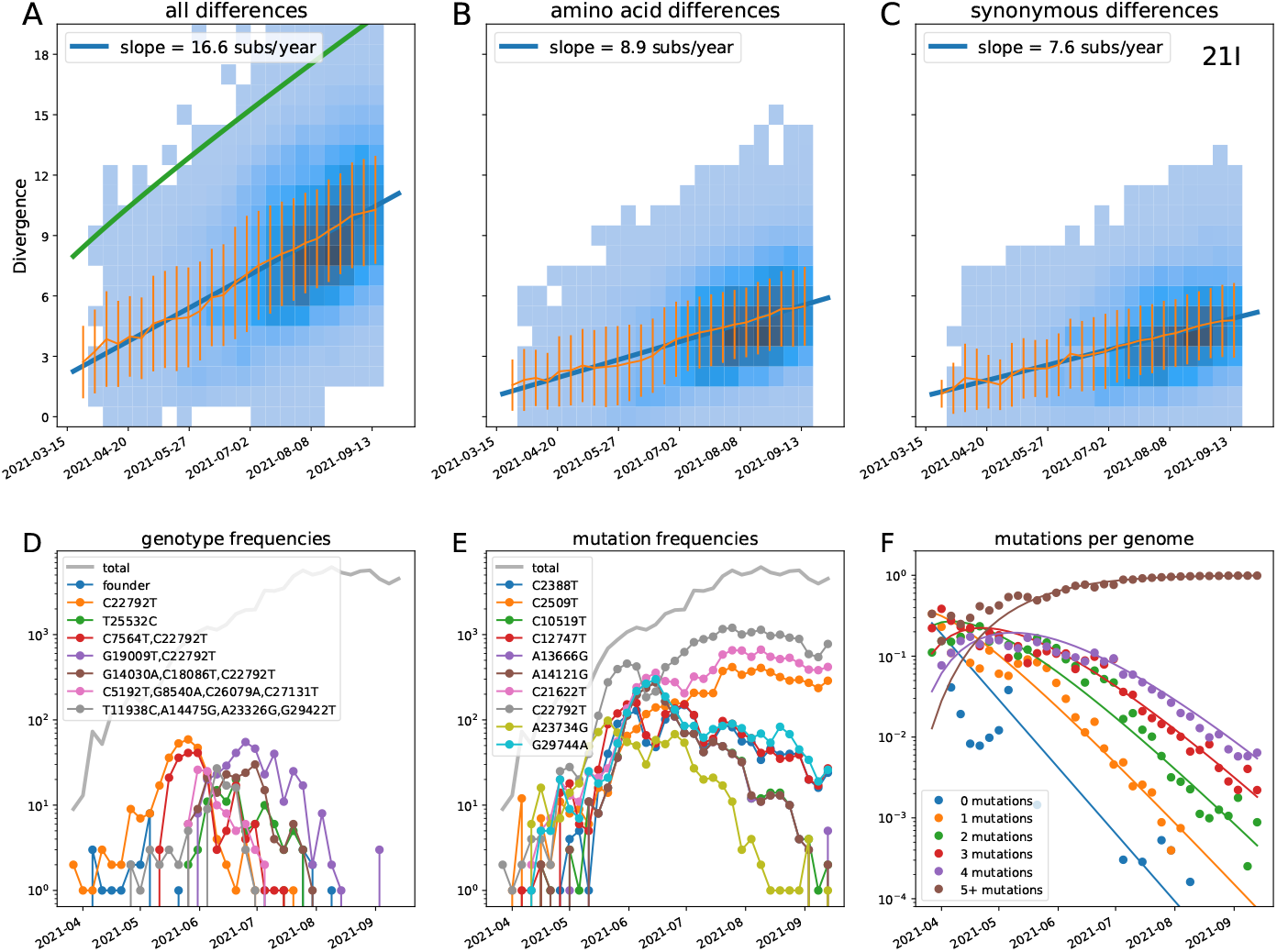
Divergence increases linearly with time in clade 21I (Delta).

**Figure S 20.**
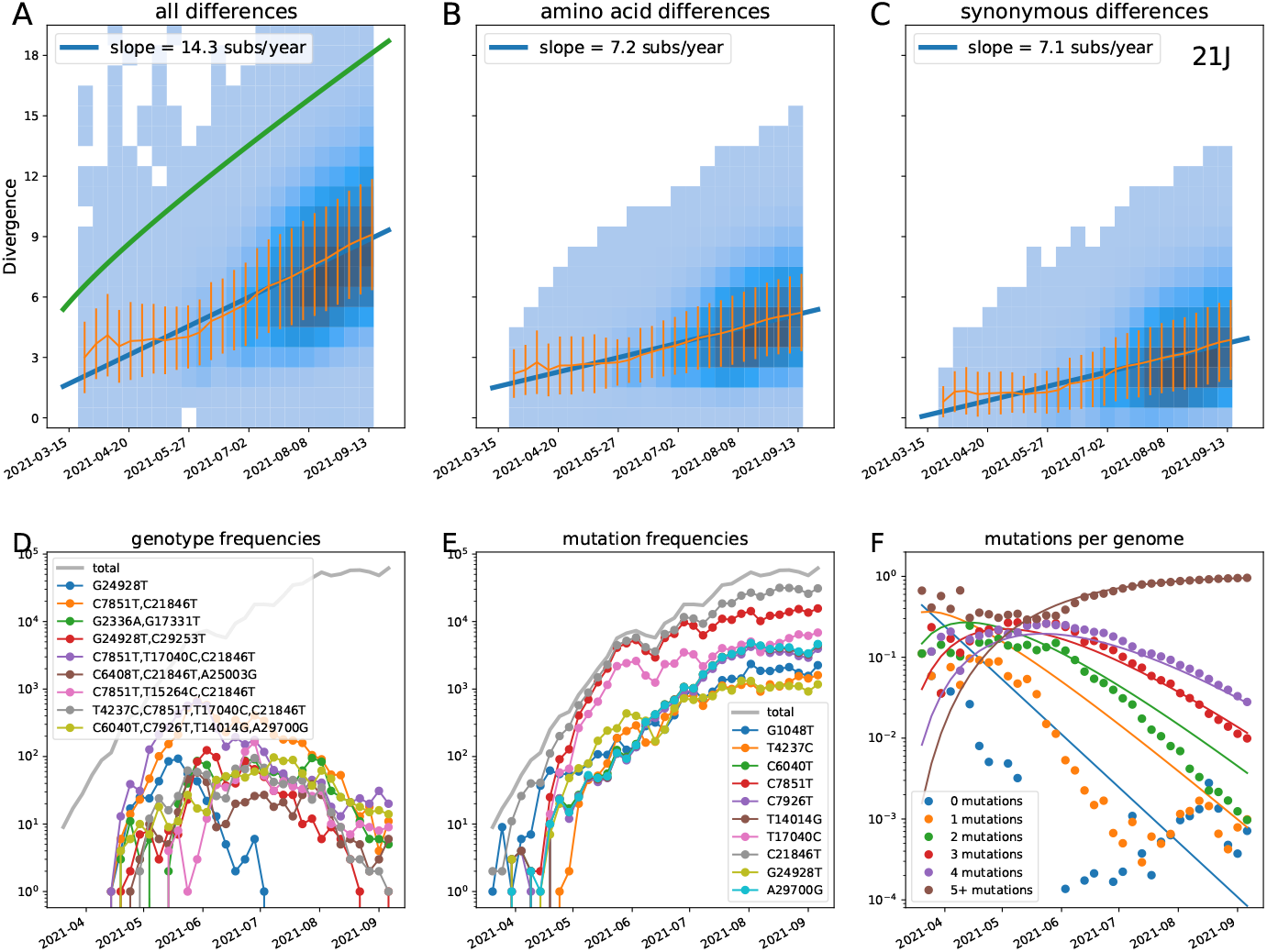
Divergence increases linearly with time in clade 21J (Delta).

**Figure S 21.**
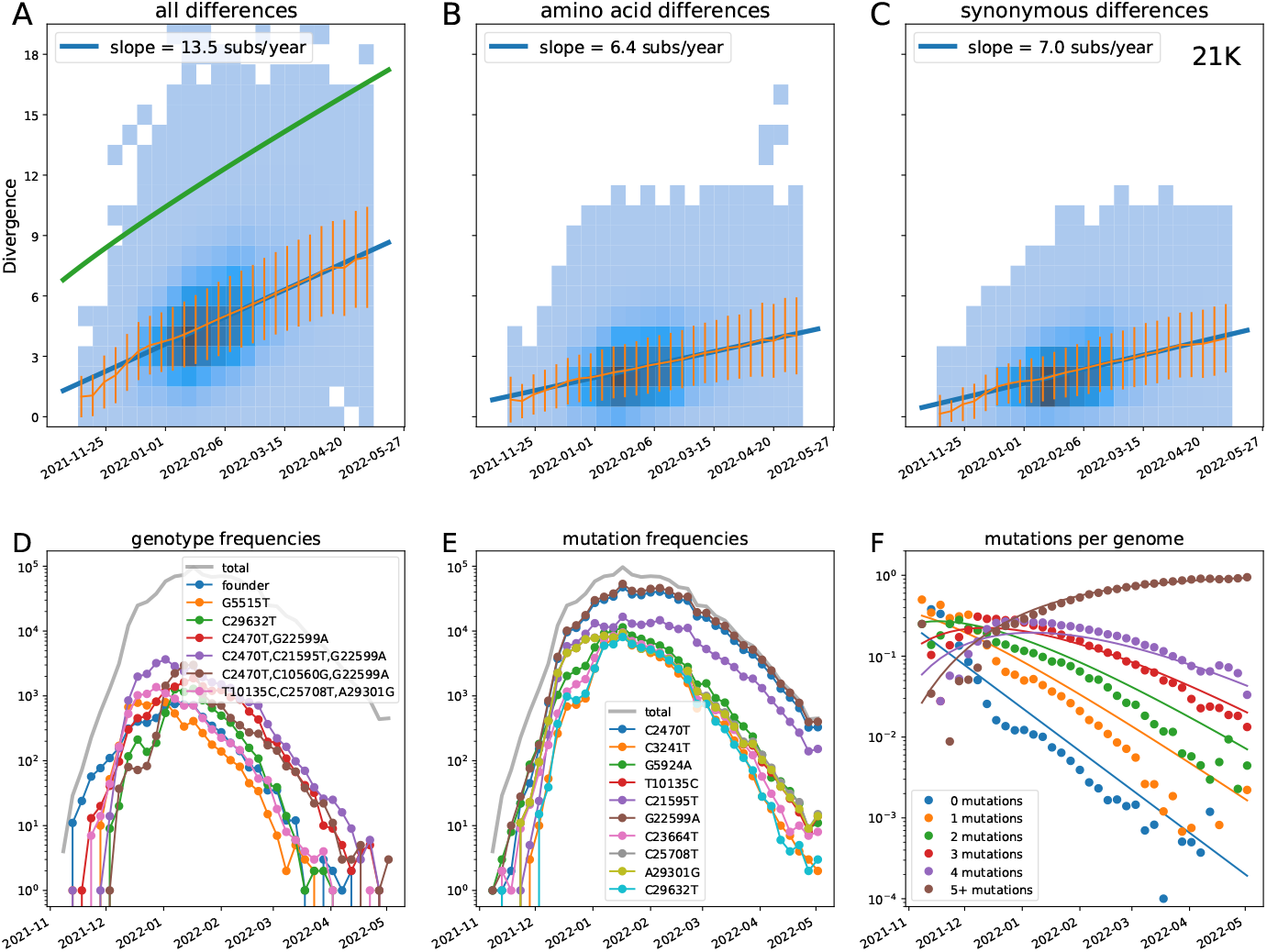
Divergence increases linearly with time in clade 21K (Omicron).

**Figure S 22.**
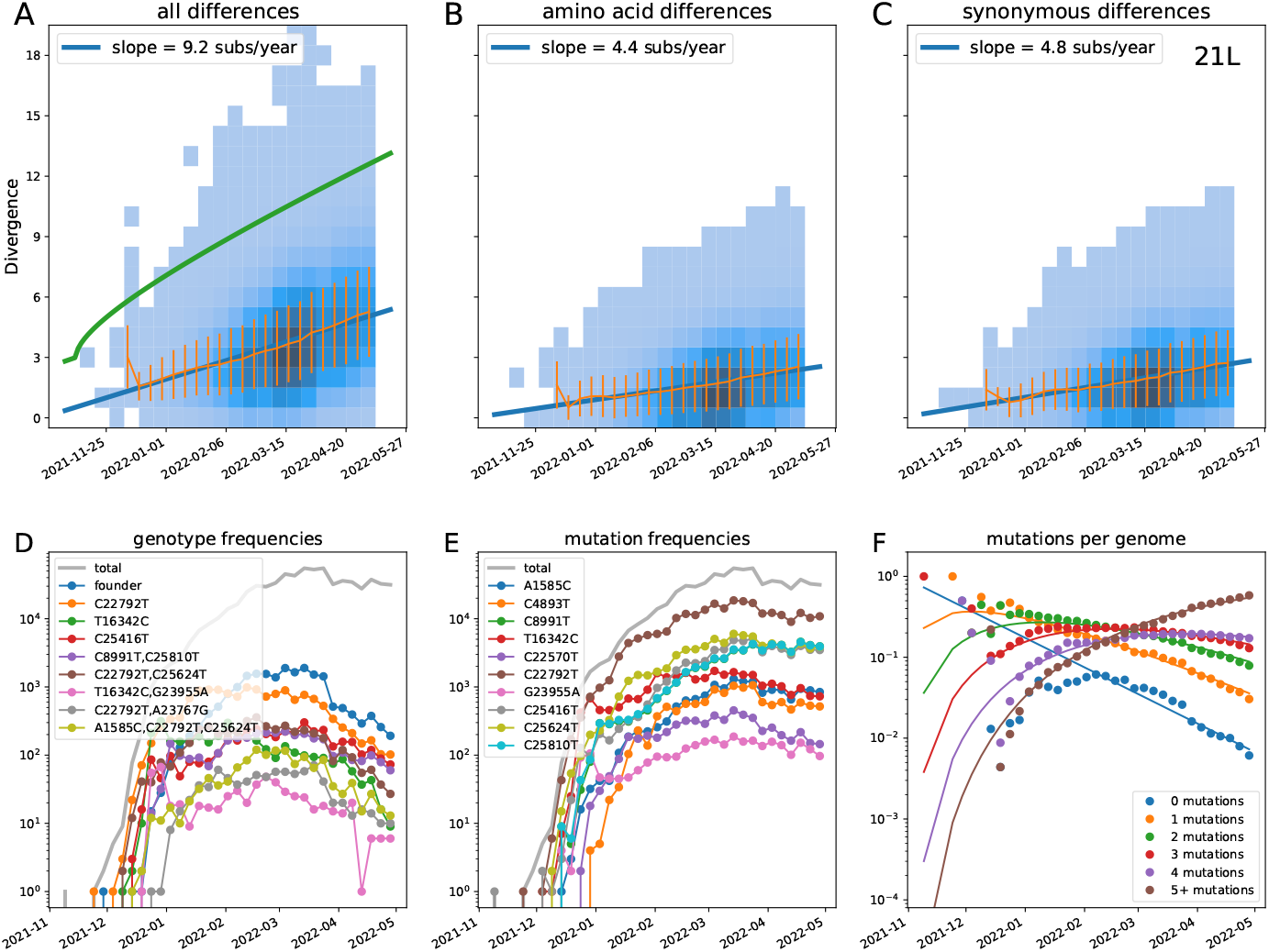
Divergence increases linearly with time in clade 21L (Omicron).

**Figure S 23.**
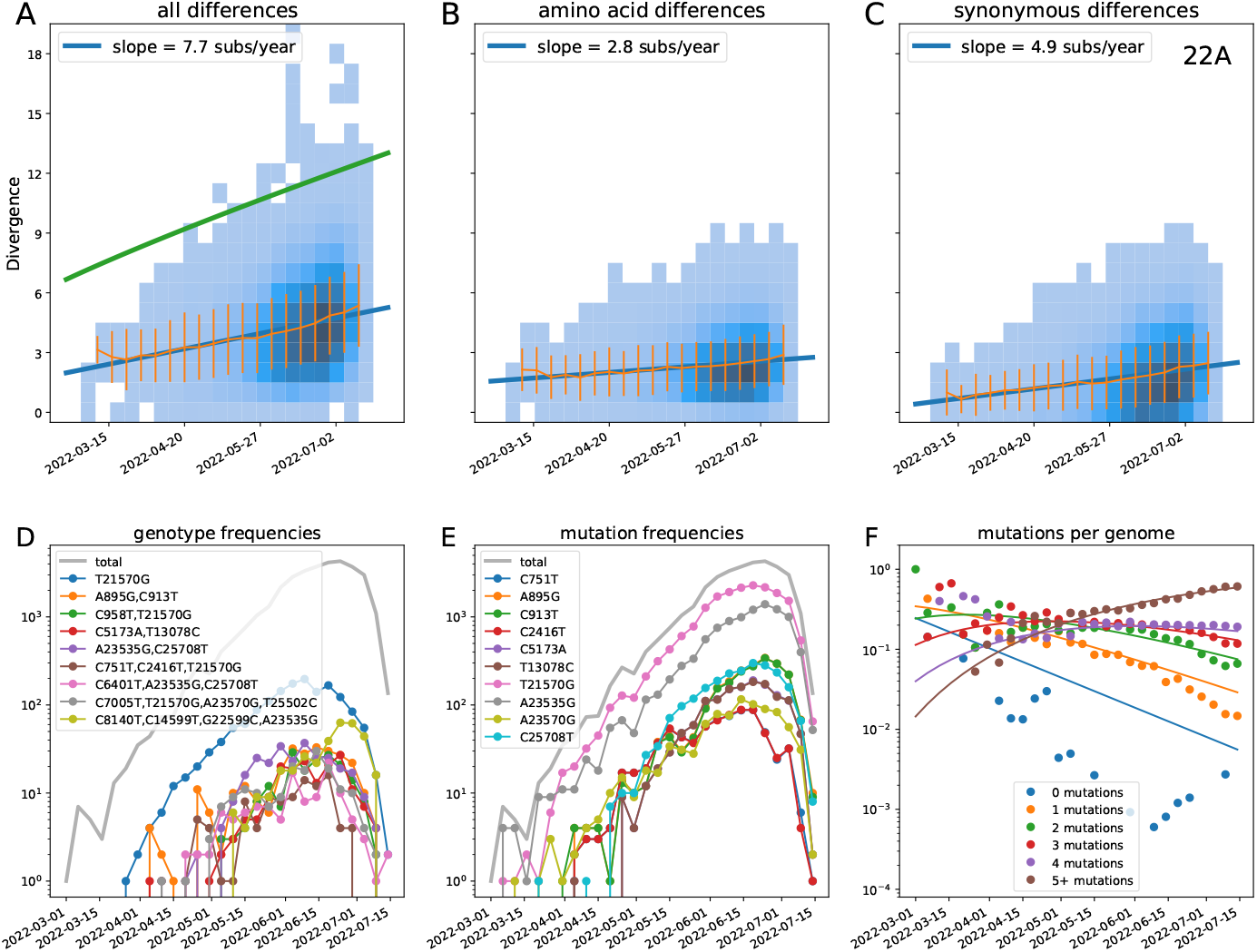
Divergence increases linearly with time in clade 22A (Omicron).

**Figure S 24.**
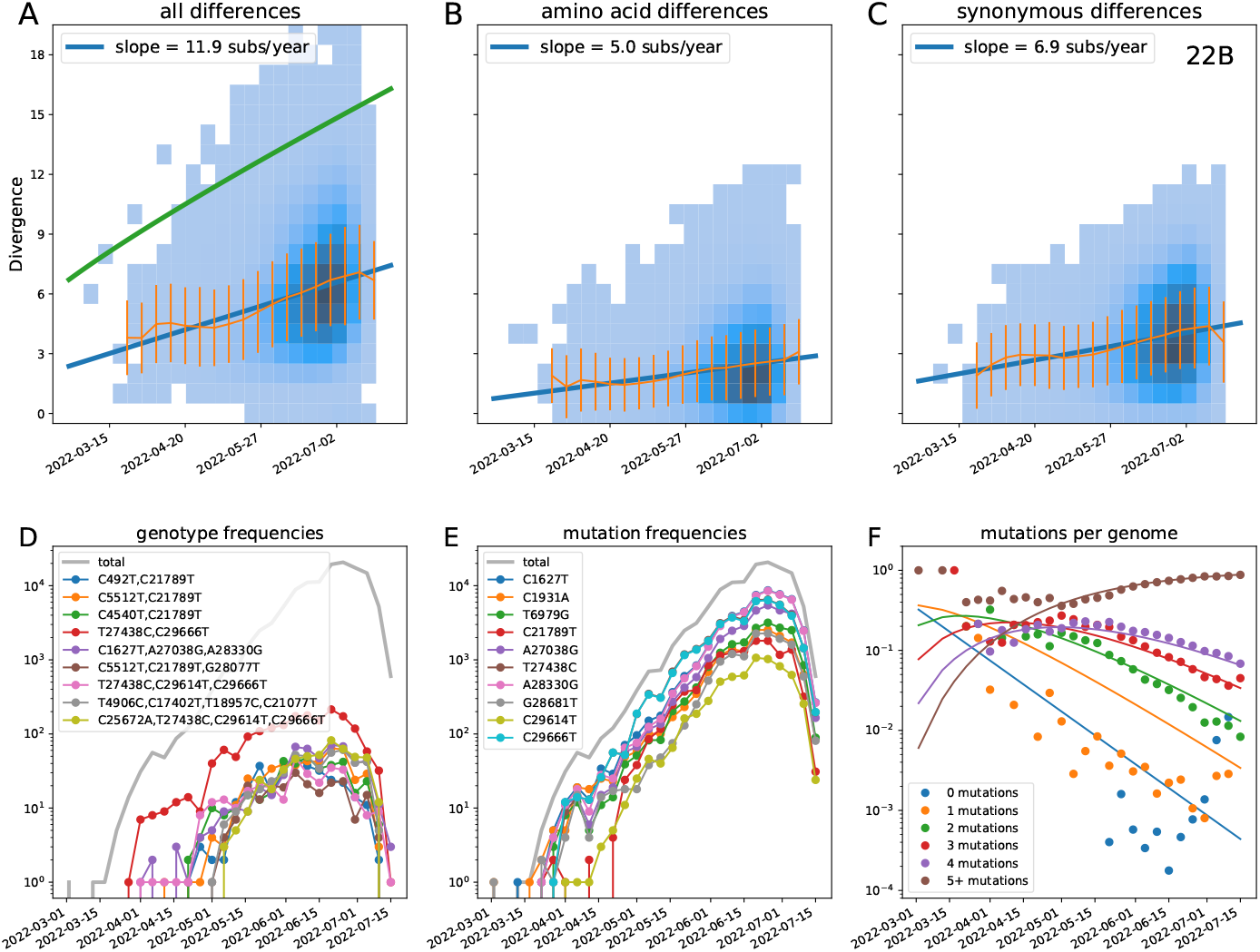
Divergence increases linearly with time in clade 22B (Omicron).

## References

Aksamentov, I., C. Roemer, E. B. Hodcroft, and R. A. Neher, 2021, Journal of Open Source Software 6(67), 3773, ISSN 2475-9066, URL https://joss.theoj.org/papers/10.21105/joss.03773.

Amicone, M., V. Borges, M. J. Alves, J. Isidro, L. Zé-Zé, S. Duarte, L. Vieira, R. Guiomar, J. P. Gomes, and Gordo, 2022, Evolution, Medicine, and Public Health 10(1), 142, ISSN 2050-6201, URL https://doi.org/10.1093/emph/eoac010.

Bhatt, S., E. C. Holmes, and O. G. Pybus, 2011, Molecular Biology and Evolution 28(9), 2443, ISSN 0737-4038, URL https://doi.org/10.1093/molbev/msr044.

Caraballo-Ortiz, M. A., S. Miura, M. Sanderford, T. Dolker, Q. Tao, S. Weaver, S. L. K. Pond, and S. Kumar, 2022, Bioinformatics 38(10), 2719, ISSN 1367-4803, URL https://doi.org/10.1093/bioinformatics/btac186.

Cele, S., F. Karim, G. Lustig, J. E. San, T. Hermanus, H. Tegally, J. Snyman, T. Moyo-Gwete, E. Wilkinson, M. Bernstein, K. Khan, S.-H. Hwa, et al., 2022, Cell Host & Microbe 30(2), 154, ISSN 1931-3128, URL https://www.sciencedirect.com/science/article/pii/S1931312822000415.

Choi, B., M. C. Choudhary, J. Regan, J. A. Sparks, R. F. Padera, X. Qiu, I. H. Solomon, H.-H. Kuo, J. Boucau, K. Bowman, U. D. Adhikari, M. L. Winkler, et al., 2020, New England Journal of Medicine 383(23), 2291, ISSN 0028-4793, publisher: Massachusetts Medical Society, URL https://www.nejm.org/doi/10.1056/NEJMc2031364.

De Maio, N., C. Walker, R. Borges, L. Weilguny, G. Slodkowicz, and N. Goldman, 2020, Issues with SARS-CoV-2 sequencing data - SARS-CoV-2 coronavirus / nCoV-2019 Genomic Epidemiology, URL https://virological.org/t/issues-with-sars-cov-2-sequencing-data/473.

Drummond, A. J., S. Y. W. Ho, M. J. Phillips, and A. Rambaut, 2006, PLOS Biology 4(5), e88, ISSN 1545-7885, publisher: Public Library of Science, URL https://journals.plos.org/plosbiology/article?id=10.1371/journal.pbio.0040088.

Elena, S. F., and R. Sanjuán, 2007, Annual Review of Ecology, Evolution, and Systematics 38, 27, ISSN 1543-592X, publisher: Annual Reviews, URL https://www.jstor.org/stable/30033851.

Faria, N. R., T. A. Mellan, C. Whittaker, I. M. Claro, D. d. S. Candido, S. Mishra, M. A. E. Crispim, F. C. S. Sales, I. Hawryluk, J. T. McCrone, R. J. G. Hulswit, L. A. M. Franco, et al., 2021, Science 372(6544), 815, publisher: American Association for the Advancement of Science, URL https://www.science.org/doi/10.1126/science.abh2644.

Ghafari, M., L. du Plessis, J. Raghwani, S. Bhatt, B. Xu, O. G. Pybus, and A. Katzourakis, 2022, Molecular Biology and Evolution 39(2), msac009, ISSN 1537-1719, URL https://doi.org/10.1093/molbev/msac009.

Ghafari, M., P. Simmonds, O. G. Pybus, and A. Katzourakis, 2021, bioRxiv, 2021.02.09.430479 Publisher: Cold Spring Harbor Laboratory Section: New Results, URL https://www.biorxiv.org/content/10.1101/2021.02.09.430479v1.

Gonzalez-Reiche, A. S., H. Alshammary, S. Schaefer, G. Patel, J. Polanco, A. A. Amoako, A. Rooker, C. Cognigni, D. Floda, A. v. d. Guchte, Z. Khalil, K. Farrugia, et al., 2022, medRxiv, 2022.05.25.22275533ISSN: 2227-5533 Type: article, URL https://www.medrxiv.org/content/10.1101/2022.05.25.22275533v1.

Hadfield, J., C. Megill, S. M. Bell, J. Huddleston, B. Potter, C. Callender, P. Sagulenko, T. Bedford, and R. A. Neher, 2018, Bioinformatics 34(23), 4121, ISSN 1367-4803, URL https://doi.org/10.1093/bioinformatics/bty407.

Hill, V., L. D. Plessis, T. P. Peacock, D. Aggarwal, R. Colquhoun, A. M. Carabelli, N. Ellaby, E. Gallagher, N. Groves, B. Jackson, J. T. McCrone, A. O’Toole, et al., 2022, The origins and molecular evolution of SARS-CoV-2 lineage B.1.1.7 in the UK, pages: 2022.03.08.481609 Section: New Results, URL https://www.biorxiv.org/content/10.1101/2022.03.08.481609v1.

Hodcroft, E. B., M. Zuber, S. Nadeau, T. G. Vaughan, K. H. D. Crawford, C. L. Althaus, M. L. Reichmuth, J. E. Bowen, A. C. Walls, D. Corti, J. D. Bloom, D. Veesler, et al., 2021, Nature, 1ISSN 1476-4687, publisher: Nature Publishing Group, URL https://www.nature.com/articles/s41586-021-03677-y.

Kemp, S. A., D. A. Collier, R. P. Datir, I. A. T. M. Ferreira, S. Gayed, A. Jahun, M. Hosmillo, C. Rees-Spear, P. Mlcochova, I. U. Lumb, D. J. Roberts, A. Chandra, et al., 2021, Nature 592(7853), 277, ISSN 1476-4687, number: 7853 Publisher: Nature Publishing Group, URL https://www.nature.com/articles/s41586-021-03291-y.

Kistler, K. E., J. Huddleston, and T. Bedford, 2022, Cell Host & Microbe 30(4), 545, ISSN 1931-3128, publisher: Elsevier, URL https://www.cell.com/cell-host-microbe/abstract/S1931-3128(22)00148-2.

Konings, F., M. D. Perkins, J. H. Kuhn, M. J. Pallen, E. J. Alm, B. N. Archer, A. Barakat, T. Bedford, J. N. Bhiman, L. Caly, L. L. Carter, A. Cullinane, et al., 2021, Nature Microbiology, ISSN 2058-5276, publisher: Nature Publishing Group, URL https://www.nature.com/articles/s41564-021-00932-w.

Korber, B., W. M. Fischer, S. Gnanakaran, H. Yoon, J. Theiler, W. Abfalterer, N. Hengartner, E. E. Giorgi, T. Bhattacharya, B. Foley, K. M. Hastie, M. D. Parker, et al., 2020, Cell 182(4), 812, ISSN 0092-8674, URL https://www.sciencedirect.com/science/article/pii/S0092867420308205.

Kryazhimskiy, S., D. P. Rice, E. R. Jerison, and M. M. Desai, 2014, Science 344(6191), 1519, publisher: American Association for the Advancement of Science, URL https://www.science.org/doi/10.1126/science.1250939.

Köster, J., and S. Rahmann, 2012, Bioinformatics 28(19), 2520, ISSN 1367-4803, URL https://doi.org/10.1093/bioinformatics/bts480.

Martin, D. P., S. Weaver, H. Tegally, J. E. San, S. D. Shank, E. Wilkinson, A. G. Lucaci, J. Giandhari, S. Naidoo, Y. Pillay, L. Singh, R. J. Lessells, et al., 2021, Cell 184(20), 5189, ISSN 0092-8674, URL https://www.sciencedirect.com/science/article/pii/S0092867421010503.

Meyer, A. G., S. J. Spielman, T. Bedford, and C. O. Wilke, 2015, Virus Evolution 1(1), vev006, ISSN 2057-1577, URL https://doi.org/10.1093/ve/vev006.

Naveca, F. G., V. Nascimento, V. C. de Souza, A. d. L. Corado, F. Nascimento, G. Silva, A. Costa, D. Duarte, K. Pessoa, M. Mejía, M. J. Brandão, M. Jesus, et al., 2021, Nature Medicine 27(7), 1230, ISSN 1546-170X, number: 7 Publisher: Nature Publishing Group, URL https://www.nature.com/articles/s41591-021-01378-7.

Neher, R. A., 2013, Annual Review of Ecology, Evolution, and Systematics 44(1), 195, eprint: 110512-135920, https://doi.org/10.1146/annurev-ecolsys- URL https://doi.org/10.1146/annurev-ecolsys-110512-135920.

Rambaut, A., E. C. Holmes, A. O’Toole, V. Hill, J. T. McCrone, C. Ruis, L. du Plessis, and O. G. Pybus, 2020, Nature Microbiology 5(11), 1403, ISSN 2058-5276, number: 11 Publisher: Nature Publishing Group, URL https://www.nature.com/articles/s41564-020-0770-5.

Shu, Y., and J. McCauley, 2017, Eurosurveillance 22(13), 30494, ISSN 1560-7917, publisher: European Centre for Disease Prevention and Control, URL https://www.eurosurveillance.org/content/10.2807/1560-7917.ES.2017.22.13.30494?crawler=true.

Strelkowa, N., and M. Lässig, 2012, Genetics 192(2), 671, ISSN 1943-2631, URL https://doi.org/10.1534/genetics.112.143396.

Tay, J. H., A. F. Porter, W. Wirth, and S. Duchene, 2022, Molecular Biology and Evolution 39(2), msac013, ISSN 1537-1719, URL https://doi.org/10.1093/molbev/msac013.

Tegally, H., M. Moir, J. Everatt, M. Giovanetti, C. Scheepers, E. Wilkinson, K. Subramoney, Z. Makatini, S. Moyo, D. G. Amoako, C. Baxter, C. L. Althaus, et al., 2022, Nature Medicine, 1ISSN 1546-170X, publisher: Nature Publishing Group, URL https://www.nature.com/articles/s41591-022-01911-2.

Tegally, H., E. Wilkinson, M. Giovanetti, A. Iranzadeh, V. Fonseca, J. Giandhari, D. Doolabh, S. Pillay, E. J. San, N. Msomi, K. Mlisana, A. von Gottberg, et al., 2021, Nature 592(7854), 438, ISSN 1476-4687.

Viana, R., S. Moyo, D. G. Amoako, H. Tegally, C. Scheepers, C. L. Althaus, U. J. Anyaneji, P. A. Bester, M. F. Boni, M. Chand, W. T. Choga, R. Colquhoun, et al., 2022, Nature 603(7902), 679, ISSN 1476-4687, number: 7902 Publisher: Nature Publishing Group, URL https://www.nature.com/articles/s41586-022-04411-y.

Volz, E., S. Mishra, M. Chand, J. C. Barrett, R. Johnson, L. Geidelberg, W. R. Hinsley, D. J. Laydon, G. Dabrera, A. O’Toole, R. Amato, M. Ragonnet-Cronin, et al., 2021, Nature 593(7858), 266, ISSN 1476-4687, number: 7858 Publisher: Nature Publishing Group, URL https://www.nature.com/articles/s41586-021-03470-x.

Wertheim, J. O., and S. L. Kosakovsky Pond, 2011, Molecular Biology and Evolution 28(12), 3355, ISSN 0737-4038, URL https://doi.org/10.1093/molbev/msr170.

Zanini, F., V. Puller, J. Brodin, J. Albert, and R. A. Neher, 2017, Virus Evolution 3(1), vex003, ISSN 2057-1577, URL https://doi.org/10.1093/ve/vex003.

Zhu, N., D. Zhang, W. Wang, X. Li, B. Yang, J. Song, X. Zhao, B. Huang, W. Shi, R. Lu, P. Niu, F. Zhan, et al., 2020, New England Journal of Medicine 382(8), 727, ISSN 0028-4793, URL https://doi.org/10.1056/NEJMoa2001017.

